# The determinants of subjective sleep depth: insights from a high-density-EEG study with serial awakenings

**DOI:** 10.1101/2021.04.06.438682

**Authors:** Aurélie Stephan, Sandro Lecci, Jacinthe Cataldi, Francesca Siclari

## Abstract

What accounts for the feeling of being deeply asleep? Standard sleep recordings only incompletely reflect subjective aspects of sleep and some individuals with so-called sleep misperception frequently feel awake although sleep recordings indicate clear-cut sleep. To identify the determinants of the subjective perception of sleep, we performed 787 awakenings in 20 good sleepers and 10 individuals with sleep misperception and interviewed them about their subjective sleep depth while they underwent high-density EEG sleep recordings. Surprisingly, in good sleepers, sleep was subjectively lightest in the first two hours of Non-rapid eye movement (NREM) sleep, generally considered the ‘deepest’ sleep, and deepest in rapid eye movement (REM) sleep. Compared to good sleepers, sleep misperceptors felt more frequently awake during sleep and reported lighter REM sleep. At the EEG level, spatially widespread high-frequency power was inversely related to subjective sleep depth in NREM sleep in both groups and in REM sleep in misperceptors. Subjective sleep depth positively correlated with dream-like qualities of reports of mental activity. These findings challenge the widely held notion that slow wave sleep best accounts for feeling deeply asleep. Instead, they indicate that subjective sleep depth is inversely related to a neurophysiological process that predominates in early NREM sleep, becomes quiescent in REM sleep and is reflected in high-frequency EEG-activity. In sleep misperceptors, this process is more frequently active, more spatially widespread, and abnormally persists into REM sleep. These findings help identify the neuromodulatory systems involved in subjective sleep depth and are relevant for studies aiming to improve subjective sleep quality.

## Introduction

What accounts for the subjective perception of being asleep? Why do we sometimes feel deeply asleep while at other times we do not? These questions may well appear trivial, as most individuals intuitively seem to know whether they were awake or asleep and how deep their sleep was. Yet the relationship between objective and subjective aspects of sleep is not as straightforward as one may think. A recent study involving 1483 participants showed for instance that conventional polysomnographic parameters only explain a small part of the variance in subjective sleep quality ^1^. Several studies have shown that most individuals tend to consistently overestimate sleep latencies ^2–5^ and conversely, appear to perceive a large part of intranight awakenings as sleep ^4,6,7^. An extreme mismatch between subjective estimations and objective measures of sleep is seen in patients with so-called ‘paradoxical insomnia’ or ‘sleep state misperception’, who feel awake during large parts of the night, although sleep recordings indicate clear-cut sleep. Despite the high prevalence of insomnia in the general population, the mechanisms underlying sleep misperception remain largely unknown ^2,8^. Determining what accounts for feeling deeply asleep is however crucial if one wants to develop targeted approaches to improve sleep quality, both in the genral population and in insomnia sufferers. Despite this obvious interest, few research studies have systematically evaluated the determinants of the subjective perception of sleep.

Sleep can be broadly divided into rapid eye movement (REM) sleep and Non-REM (NREM) sleep. REM sleep is characterized by rapid eye movements and fast, low-amplitude electroencephalographic (EEG) activity, whereas NREM sleep features high-amplitude, low-frequency EEG slow waves and graphoelements called spindles. According to the current classification of sleep stages, NREM sleep can be further divided into three stages, in which the EEG becomes progressively slower (stages NREM 1, 2 and 3). Stage NREM 3, in which high-amplitude slow waves occur nearly continuously, is referred to as ‘slow wave sleep’. This stage is also called ‘deep’ sleep, presumably because early studies showed that arousal thresholds were high in this stage and varied as a function of slow wave activity (SWA) ^9^. But is objectively defined ‘deep’ sleep necessarily perceived as subjectively deep?

Early work in which awakenings were performed serially in the falling asleep period revealed that approximately 50% of subjects felt asleep 4 min after the first sleep spindle and 90% after 16 min. However, subsequent studies performing awakenings in stable NREM stage 2 sleep at different times of the night revealed that subjects felt awake during polysomnographically defined sleep in a high proportion of cases ^3,5,10–13^ (reaching 40-45% in some studies ^5,14,15^), suggesting a dissociation between standard sleep parameters and the subjective estimation of sleep. Studies sampling sleep perception in REM sleep reported variable, although overall lower objective-subjective discrepancies compared to NREM sleep stage 2 ^5,11,13,16^.

More recent studies based on retrospective estimations of sleep duration given upon awakening in the morning reported positive correlations between the degree of sleep underestimation and some EEG features, including alpha ^17^, sigma ^17^ and/or beta power ^17,18^, the delta/beta power ratio ^19^, delta-alpha sleep patterns ^20^ and signatures of ‘light’ sleep within ‘deep’ sleep ^21^. One study combining serial awakenings with EEG spectral power measures found that feeling awake instances in light NREM sleep stages 1 and 2 sleep were associated with higher relative alpha, beta and gamma power compared to instances of feeling asleep ^22^.

In the present study we aimed to probe the moment-to-moment perception of sleep in a large number of instances, across different sleep stages, while recording brain activity with high-density EEG, which combines the excellent temporal resolution of EEG with an increased spatial resolution upon source modeling. This technique has been used successfully in the past to identify local correlates of subjective experiences during sleep, including dream contents ^23^. Studies using techniques with a high spatial resolution to record brain activity, including positron emission tomography ^24^, functional magnetic resonance ^22^ and high-density EEG ^25^ have indeed reported localized changes in brain activity in insomnia patients, opening the possibility that they may play a role in sleep perception. To include a broad range of different levels of sleep perception, we performed the same paradigm in 20 good sleepers, as well as in 10 insomnia sufferers who severely underestimated their sleep duration with respect to the general population (sleep misperceptors) ^4^.

## Results

Our protocol comprised an undisturbed baseline high-density EEG sleep recording followed by two recordings during which participants were awakened multiple times with a computerized alarm sound ^26^ (see star methods for details). Awakenings were scheduled after 10 minutes of consolidated NREM or REM sleep. After each awakening, participants were asked to estimate whether they had been feeling awake (FAW) or asleep (FAS) prior to the alarm sound. When they had felt asleep, they also had to estimate the degree of sleep depth on a scale from 1 to 5. In addition, they had to describe the last thing going through their mind before the alarm sound and to rate specific cognitive aspects of their experience on a 1-5 point scale ^23,26^ (see star methods for more details).

Overall, we performed 787 awakenings. Good sleepers and sleep misperceptors did not differ in the number of awakenings per subject [26.7 +/- 7.1 vs. 25.3 +/- 5.7, t-test p=0.56], the proportion of awakenings in NREM (stages N2& N3) vs. REM sleep [NREM good sleepers: 76.8 +/- 4.6%; sleep misperceptors: 75.8+/- 2.8%, t-test p=0.42] or the time spent asleep before each awakening [NREM: 13.7 +/- 1.2 min vs 14.9 +/- 1.8 min, t-test p=0.07; REM: 21.6 +/- 7.9 min vs 20.1 +/- 3.8 min, t-test p=0.49]. More exhaustive information on the number of trials in different conditions can be found in Figure S1.

### Effect of serial awakenings

As expected, during the serial awakening nights, sleep was more fragmented compared to the undisturbed baseline night. A mixed model analysis revealed an effect of the type of night on sleep efficiency, total sleep time, the proportion of NREM stages 1 and 2 and REM sleep, as well as REM latency (Table 1). Importantly, removing group (misperceptors vs. good sleepers) as a random factor from the model did not change the results, indicating that serial awakenings had similar effects on sleep structure in both groups. The two serial awakening nights were comparable between groups in terms of sleep parameters and proportion of FAW instances (linear mixed model revealed no effect of night and removing group as random effect did not affect results).

**Table 1.** Table of sleep parameters in baseline and serial awakening nights. Sleep parameters for baseline and serial awakening nights. Results of mixed model analyses explaining the sleep parameter by the type of night (baseline versus serial awakening), including group as a random factor. P-values (corrected and uncorrected for multiple comparisons) are shown. Bold font indicates parameters that significantly differed between baseline and serial awakening nights.

|  | Baseline | Serial Awakening | Chi-square | P-value | P corrected |
| --- | --- | --- | --- | --- | --- |
| Total recording time (min) | 423 ± 29 | 413 ± 31 | 2,34 | 0,13 | 1 |
| **Total sleep time (TST, in min)** | **372 ± 42** | **317 ± 52** | **26,74** | **< 0,001** | **< 0,001** |
| **Wake after sleep onset (min)** | **45 ± 33** | **89 ± 39** | **29,31** | **< 0,001** | **< 0,001** |
| **Sleep efficiency (%)** | **89 ± 8.1** | **78 ± 9.8** | **27** | **< 0,001** | **< 0,001** |
| Sleep latency (min to N1) | 6.6 ± 4.6 | 5.8 ± 5.2 | 0,48 | 0,48 | 1 |
| **REM latency (min)** | **86 ± 31** | **132 ± 63** | **13,5** | **< 0,001** | **0,004** |
| **N1 (% of TST)** | **4.3 ± 2.6** | **7.6 ± 4.2** | **15,4** | **< 0,001** | **0,002** |
| **N2 (% of TST)** | **52 ± 5.9** | **58 ± 6.6** | **15,7** | **< 0,001** | **0,001** |
| N3 (% of TST) | 21 ± 5.8 | 19 ± 6.7 | 3,06 | 0,08 | 1 |
| **REM (% of TST)** | **22 ± 4.6** | **26 ± 4.3** | **43,6** | **< 0,001** | **< 0,001** |

**Table 2.** List of generalized and linear mixed models. All models referred to in the result section are summarized in this table, with specifications and reference to the figure in which their result are displayed. QA = Qualitative aspect = thought-like, perceptual, Richness and complexity, Duration, Control, Lucidity; Degree = degree of subjective sleep depth; Event = FAW vs FAS; time = time since lights off; time_asleep = time spent asleep before each awakening.

|  | Formula | Details | Figure |
| --- | --- | --- | --- |
| #1 | Perceived State ~ Group*Stage + (1\|sub) + (1\|time_asleep) | Stage = NREM vs REM | Fig.1A |
| #2 | Perceived State ~ Group*Time + (1\|sub) + (1\|time_asleep) | In NREM only as FAW are quasi inexistent in REM | Fig.1B |
| #3 | Degree ~ Group*Stage*Time + (1\|sub) + (1\|time_asleep) | Stage = NREM vs REM, FAS only | Fig.1C-D |
| #4 | CE_presence ~ Group*Stage | CE_presence = CE vs NE, FAS only | Fig.2A |
| #5 | QA ~ Group*Stage + (1\|sub) + (1\|time_asleep) | One model per QA, FAS only | Fig.2B |
| #6 | Degree ~ Group*Stage*CE + (1\|sub) + (1\|time_asleep) | One mode per QA, FAS only |  |
| #7 | Degree ~ Group*Stage*QA + (1\|sub) + (1\|time_asleep) | One mode per QA, FAS only |  |
| #8 | Degree ~ PSD*Group + (1\|sub) + (1\|time) | One model per electrode per frequency band in NREM and REM separately | Fig.3<br>Fig.5 |
| #9 | PSD ~ Time*Group + (1\|sub) + (1\|time_asleep) | One model per frequency band in NREM, averaged over all electrodes | Fig. S.4 |
| #10 | Degree ~ SP*Group + (1\|sub) + (1\|time) | One model per electrode per spindle parameter in NREM | Fig.4 |

Next, we evaluated how consistent ratings of sleep perception were among participants. Sleep misperceptors, but not good sleepers, demonstrated a significant correlation between the proportion of FAW in the serial awakening nights and the sleep perception index of the baseline night (ratio between subjective and objective total sleep time in percent) *[r2 = −0.78, p = 0.003 vs r2 = −0.40 p>0.05]*; as well as between the proportion of FAW in the two serial awakening nights *[r2 = 0.85, p < 0.001 vs r2 = 0.40, p>0.05;* **Figure S.2B**].

### Effect of group, sleep stage and time of the night on perceived state (FAW/FAS)

To evaluate whether the proportion of FAS and FAW differed between sleep misperceptors and good sleepers, REM and NREM (stages N2 & N3) sleep, and according to the time of the night, we computed a generalized linear mixed model, with perceived state (FAW vs. FAS) as the categorical dependent variable, group (good sleepers vs. sleep misperceptors), stage (NREM vs. REM) and/or time (hours after lights off) as fixed factors, and subject and time spent asleep ^27^ as random factors (#1, #2).

As expected, sleep misperceptors felt more often awake than good sleepers *[main effect of group, χ^2^(1) = 11.0, p=0.001, **Figure 1A**, estimates and post-hoc results in **Table S.1B-C**]*. Both groups felt more often awake in NREM than REM sleep *[main effect of stage: χ^2^(1) = 7.7, p=0.005, **Figure 1A, Table S.1B-C,** no group-stage interaction, NREM: 12.3% and 39.6% in good sleepers and sleep misperceptors respectively, REM: 0.8% and 9.8%; χ^2^(1) = 0.28, p=0.59, **Table S.1A**]*.

**Figure 1.**
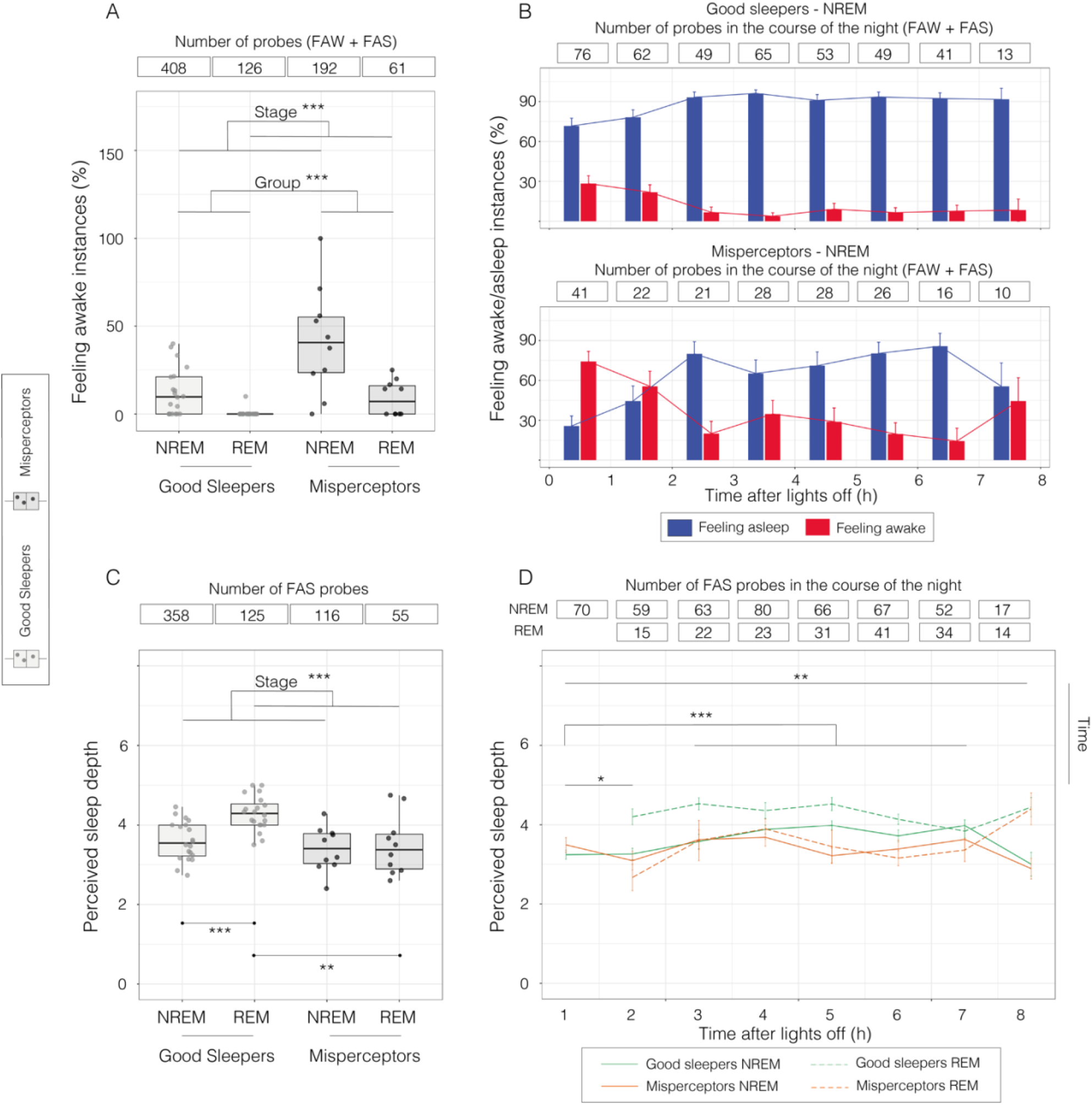
Determinants of sleep perception. Perceived state (feeling awake vs feeling asleep; A, B) and degree of perceived sleep depth (scale from 1 to 5; C, D) as a function of group (good sleepers/misperceptors), stage (REM/NREM) and time of the night. **A.** Proportion of feeling awake instances in good sleepers (n=20, grey) and misperceptors (n=10, black) in NREM and REM sleep. **B.** Average proportion of feeling awake (red) and feeling asleep instances (blue) +/- standard error across trials in good sleepers (upper row) and misperceptors (bottom row) in NREM sleep in the course of the night. **C.** Degree of perceived sleep depth in good sleepers (grey) and misperceptors (black) in NREM and REM sleep (1=minimal; 5= maximal). **D.** Average degree of perceived sleep depth +/- standard error across trials in good sleepers (green) and misperceptors (orange) in NREM (solid line) and REM (dashed line) in the course of the night. In boxplot figures (A, C), each point represents an individual, the box displays the 25^th^ to 75^th^ percentile of data and the vertical bars shows the confidential interval. Post-hoc of significant main and interaction effects evaluated with generalized linear mixed models are indicated with lines and asterisks indicating the level of significance: *\* p < 0.05, **p < 0.01, ***p < 0.001*, including probability of FAW/FAS as a function of group and stage (A); group and time of night (B). Degree of perceived sleep depth as a function of group, stage and time of night (C, D). All models included subject identity and time spent asleep as random factors. NREM refers to stages N2 and N3.

Within NREM sleep, the proportion of FAW changed significantly in the course of the night in both groups *[main effect of time of night: χ^2^(7) = 48.1, p<0.0001, **Figure 1B**, no group-time interaction effect, **Table S.2A**]*, with most FAW instances occurring in the first two, followed by the last hour of the night (**Table S.2B-C**). Characteristics of FAW in REM sleep were not further analyzed as they were rare in this stage.

### Effect of group, sleep stage and time of the night on perceived sleep depth

We then evaluated, for all FAS instances, how perceived sleep depth differed between groups, according to sleep stage and time of night. Here, we performed a linear mixed model, with perceived sleep depth (i.e. degree, from 1 to 5, 5=deepest) as the ordinal dependent variable, and the same fixed and random factors as in the model described above (#3). Good sleepers felt more deeply asleep in REM than in NREM sleep, while sleep misperceptors reported equivalent subjective sleep depths in REM and NREM sleep *[significant group - sleep stage interaction effect: χ^2^(1) 4.2, p=0.041, **Table S.3B-C**]*. Similar to FAW instances, subjective sleep depth within NREM sleep, but not REM sleep markedly and consistently changed in the course of the night – with subjects feeling most lightly asleep in the first two hours, followed by the last hour of the night *[main effect of time of night, χ^2^(7) = 59.1, p <0.0001, interaction effect between time and stage χ^2^(6)=13.2, p=0.04, **Figure 1D**, **Table S.3B-C**]*.

### Additional verifications

Perceived state and sleep depth were comparable between stages N2 and N3 but differed from REM sleep in both cases *[main effect of group: perceived state: χ^2^(1)=16.82, p<0.0001 and of stage: perceived state: χ^2^(1)=12.37, p=0.002, **Table S.4B-C**; degree of perceived sleep depth: χ^2^(1)=9.32, p=0.01, **Table S.5B-C**, **Figure S.3**]*. For all further analyses, stages N2 and N3 were therefore analyzed together (as NREM). All models evaluating the effect of time since lights off (#2, #3) were also performed with “time since first sleep onset”, defined as the time in hours since the first page of N1; results remained unchanged (Table S.6B-C and 7B-C).

### Perceived sleep depth and conscious experiences

In line with previous serial awakening paradigms investigating conscious experiences in sleep ^26,28^, both good sleepers and sleep misperceptors were more likely to report conscious experiences (CE) when feeling asleep in REM sleep compared to NREM sleep (#4) *[main effect of stage, χ^2^(1) = 12.6; p=0.0004, **Figure 2.A, Table S.8B-C**, no effect of group; no interaction effect between group and stage]*. In both groups, experiences reported upon awakening from REM sleep were more perceptual, richer, longer and associated with lower degrees of lucidity compared to NREM experiences (#5) *[significant main effect of stage on perception χ^2^(1) = 32.01; p<0.0001, **Table S.10B-C**; richness & complexity χ^2^(1) = 10.58, p=0.001, **Table S.11B-C**; duration χ^2^(1) = 28.23, p<0.0001, **Table S.12B-C**; lucidity χ^2^(1) =5.33, p=0.02,, **Table S.14B-C**; **Figure 2.B**, no group-stage interaction effects]*. Regardless of sleep stage, good sleepers and sleep misperceptors differed only in the thinking dimension, with sleep misperceptors reporting significantly more thought-like experiences compared to good sleepers *[main effect of group, χ^2^(1) = 9.01, p=0.003, **Table S.9B-C**]*. A qualitative analysis of conscious experiences revealed that sleep misperceptors referred to their sleep and worried about not being able to fall asleep in 13.8 % of cases, while none of the good sleepers did. Examples of experiences associated with FAS and FAW reports can be found in Annex S.1.

**Figure 2.**
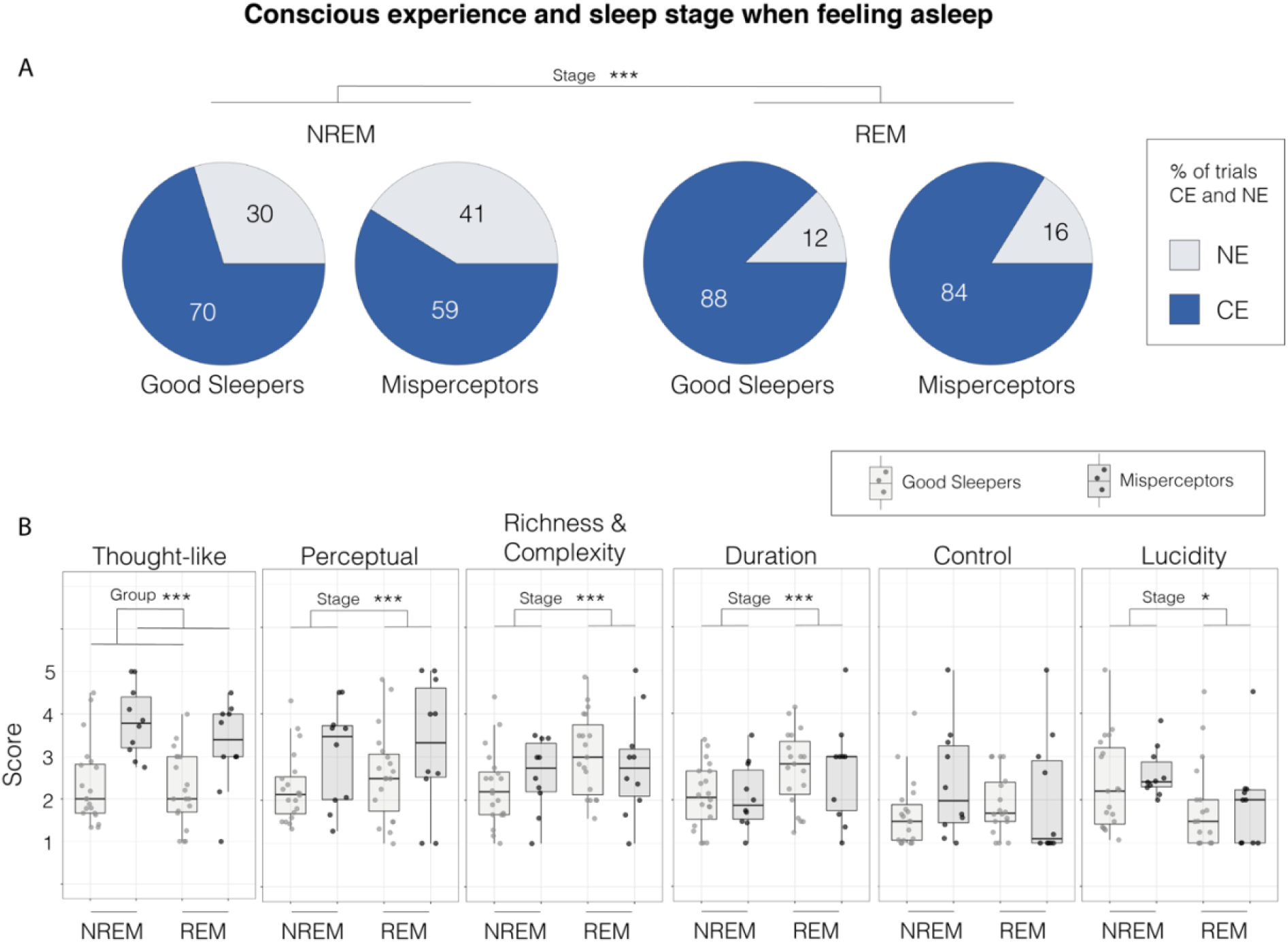
Conscious experiences in sleep and sleep perception. **A.** Probability of presence (CE) and absence (NE) of conscious experience when feeling asleep. **B.** Average scores for different scales evaluating qualitative aspects of experiences, shown for good sleepers (grey) and misperceptors (black), in NREM and REM sleep, when feeling asleep (1=minimal; 5= maximal). Each point represents an individual, the box displays the interquartile range and the vertical bars show the confidential interval. Post-hoc of significant main and interaction effects evaluated with generalized linear mixed models are indicated with lines and asterisks indicating the level of significance: *\* p < 0.05, **p < 0.01, ***p < 0.001*. (A) probability of CE/NE and (B) CE content score as a function of group and stage. All models included subject identity and time spent asleep as random factors. NREM refers to stages N2 and N3.

The presence or absence of conscious experiences did not predict subjective sleep depth (#6) *[main effect of C.Exp, χ^2^(1) = 2.66, p>0.05; no interaction effect between C.Exp and group, χ^2^(2) = 2.06, p>0.05; no interaction C.Exp and stage effect, **Table S.15A-B**]*. However, the more good sleepers and sleep misperceptors felt asleep, the richer, more complex and perceptual the experience, regardless of sleep stage (#7) *[main effect of perception, χ^2^(4)=19.8, p=0.0006, **Table S.17B-C**; and richness & complexity, χ^2^(4)=17.86, p=0.001, **Table S.18B-C**; no interaction effects with group and/or stage]*.

### EEG activity and perceived sleep depth in NREM sleep

To determine how absolute EEG spectral power in different frequency bands related to perceived sleep depth in NREM sleep, we computed a linear mixed model per frequency band and electrode, including power spectral density and group as fixed factors and subject identity and time of night as random factors (#8). We found that sigma power (12-16 Hz) in good sleepers and beta power (18-29.5 Hz) in both groups significantly predicted lower subjective sleep depth (**Figure 3A, first two rows**). No interaction effect between spectral power and groups was found (**Figure 3A, last row**), suggesting a similar relationship between power in these frequency bands and sleep perception in both groups. To map the topography of the effects in the sigma and beta power range on sleep depth, we performed the same analysis on source reconstructed data (**Figure 3B**). We found significant effects of sigma power (in good sleepers) and beta power (in sleep misperceptors) on subjective sleep depth, which were spatially widespread, peaking in central brain regions, and sparing only small portions of the frontopolar, inferior and medial temporo-occipital cortices. For beta power, sleep misperceptors displayed more widespread effects compared to good sleepers, extending to fronto-polar and occipital areas, although this difference between groups did not resist our stringent correction for multiple comparisons.

**Figure 3.**
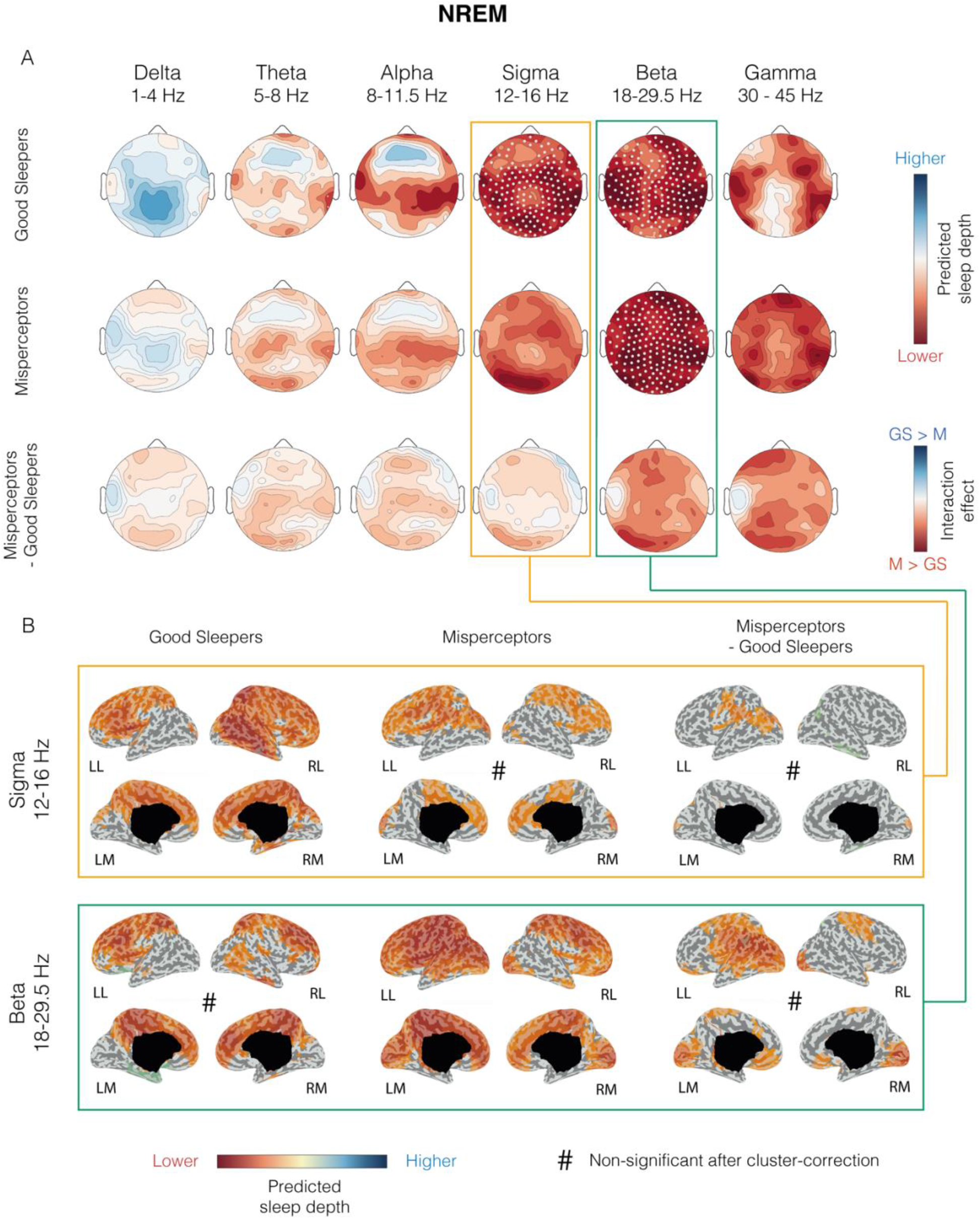
Absolute power spectral density and perceived sleep depth in NREM sleep. **A.** Results of linear mixed models explaining perceived sleep depth by power spectral density in good sleepers (n=20, top row) and misperceptors (n=10; middle row). Group interaction effect is displayed in bottom row. Absolute power spectral density was averaged over the 120s preceding the awakening. The effect of each frequency band was evaluated in separate models. All models included subject identity and time of night as random factors. Wald statistics values (squared ratio of the fixed factor estimate over its standard error) are shown at the scalp level for each electrode; electrodes with a significant effect after a cluster-and probability-based correction for multiple comparisons (p<0.05) are marked white. **B.** Cortical distribution of Wald statistics at the source level for selected frequency bands (based on results shown in A). The same procedure described in A was used. Voxels with non-significant results are colored in grey. Contrasts that did not survive the correction for multiple comparisons are marked with a hash sign. NREM refers to stages N2 and N3. *LL = left lateral, RL = right lateral, LM = left medial, RM = right medial*.

To determine how our spectral power findings on subjective sleep depth related to the course of subjective sleep depth across a night of sleep, we investigated how spectral power in different frequency bands changed during the night (#9, **Figure S.4**). All frequency bands showed significant changes in the course of the night in NREM sleep in both good sleepers and sleep misperceptors (**Table S.22-27**). In the first hour after lights off, delta power was significantly lower in sleep misperceptors than in good sleepers (**Table S.22D**). However, despite the serial awakenings, delta power significantly decreased in both groups across the night, as would be expected for an undisturbed night of sleep **(Table S.22B-D)**. Only beta power inversely mirrored the course of subjective sleep depth in both groups, decreasing early in the night and increasing towards the end of the night (**Table S.26B-C**), consistent with our findings relating beta power to lower subjective sleep depth in both groups (Fig. 3A).

Since sigma power is known to reflect sleep spindle activity and predicted lower sleep depth in good sleepers, we performed a spindle detection analysis and examined how different spindle parameters related to sleep depth (#10). We found that higher spindle amplitude, but not the density of spindles, predicted a lower perceived sleep depth (**Figure 4**) in both groups. In good sleepers, a higher fast spindle density also predicted lower sleep depth. Similar to sigma power findings, these effects were spatially widespread.

**Figure 4.**
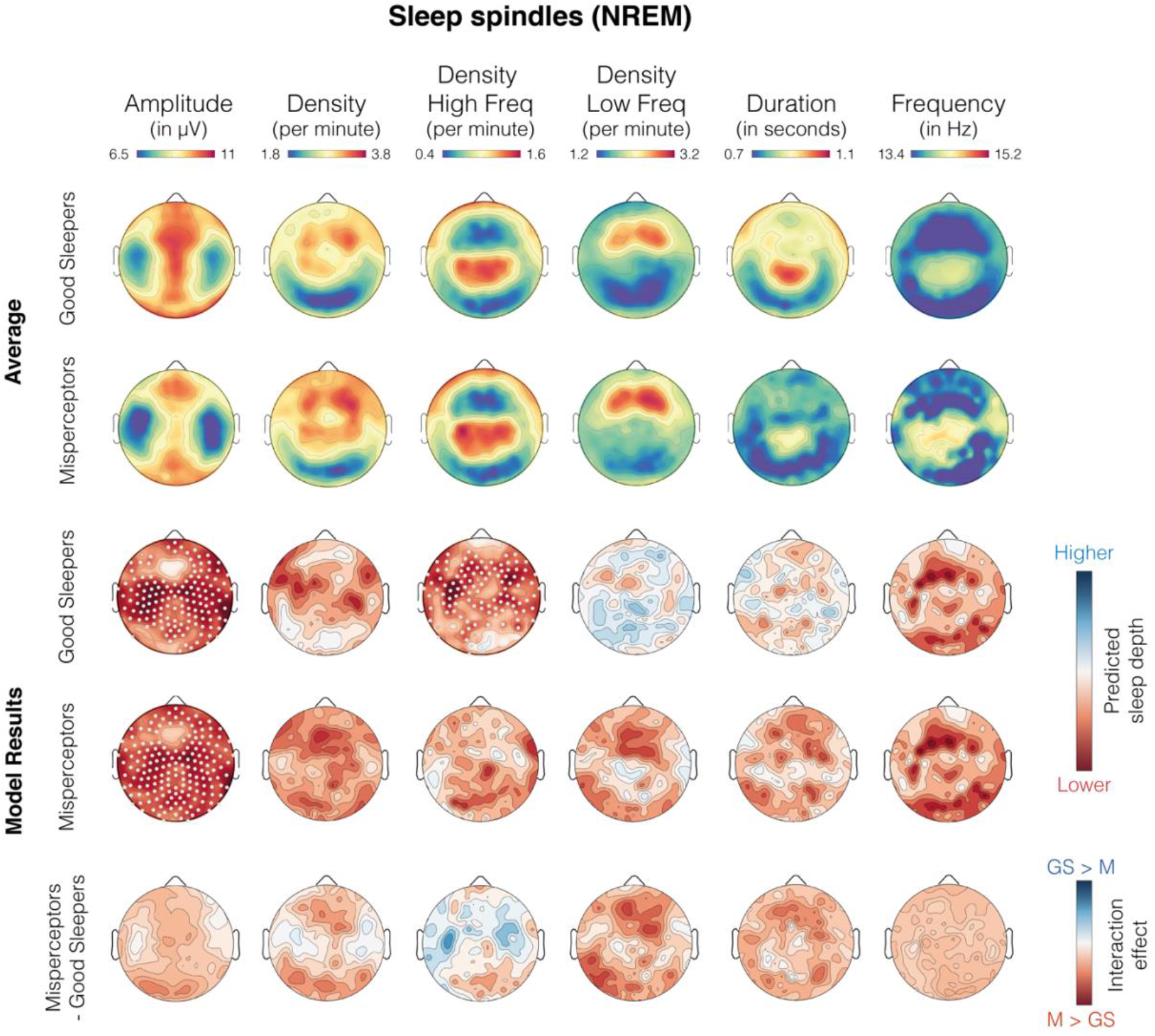
Spindles and perceived sleep depth in NREM sleep. Topographic distribution of average spindle parameters at the scalp level in good sleepers (n=20, top row) and sleep misperceptors (n=10, second row). Results of linear mixed models explaining the degree of perceived sleep depth by spindle parameters in good sleepers (third row), misperceptors (fourtth row), and group interaction effect (bottom row). Spindle parameters were extracted and averaged over the 120s preceding the awakening. The effect of each parameter was evaluated in separate models. All models included subject identity and time of night as random factors. Wald statistics values (squared ratio of the fixed factor estimate over its standard error) are shown at the scalp level for each electrode; electrodes with a significant effect after a cluster-and probability-based correction for multiple comparisons (p<0.05) are marked white. NREM refers to stage N2 and N3.

### EEG activity and perceived sleep depth in REM sleep

In REM sleep, beta (18-29.5 Hz) and gamma power (30-45 Hz) significantly predicted lower subjective sleep depth in sleep misperceptors (**Figure 5A**). A non-significant negative correlation was also seen for all other frequency bands in sleep misperceptors. In good sleepers, no significant effects were found, although a weak and non-significant tendency for alpha, sigma and beta power to predict lower sleep depth was seen. Models explaining perceived sleep depth with source reconstructed EEG activity showed again a widespread effect of beta and gamma on perceived sleep depth in sleep misperceptors, sparing only small islands of the lateral temporal and fronto-polar cortices (**Figure 5B**). As expected, corrected group interaction effects suggested that the predictive power of beta and gamma activity was significantly higher in sleep misperceptors than good sleepers (**Figure 5B**, last column).

**Figure 5.**
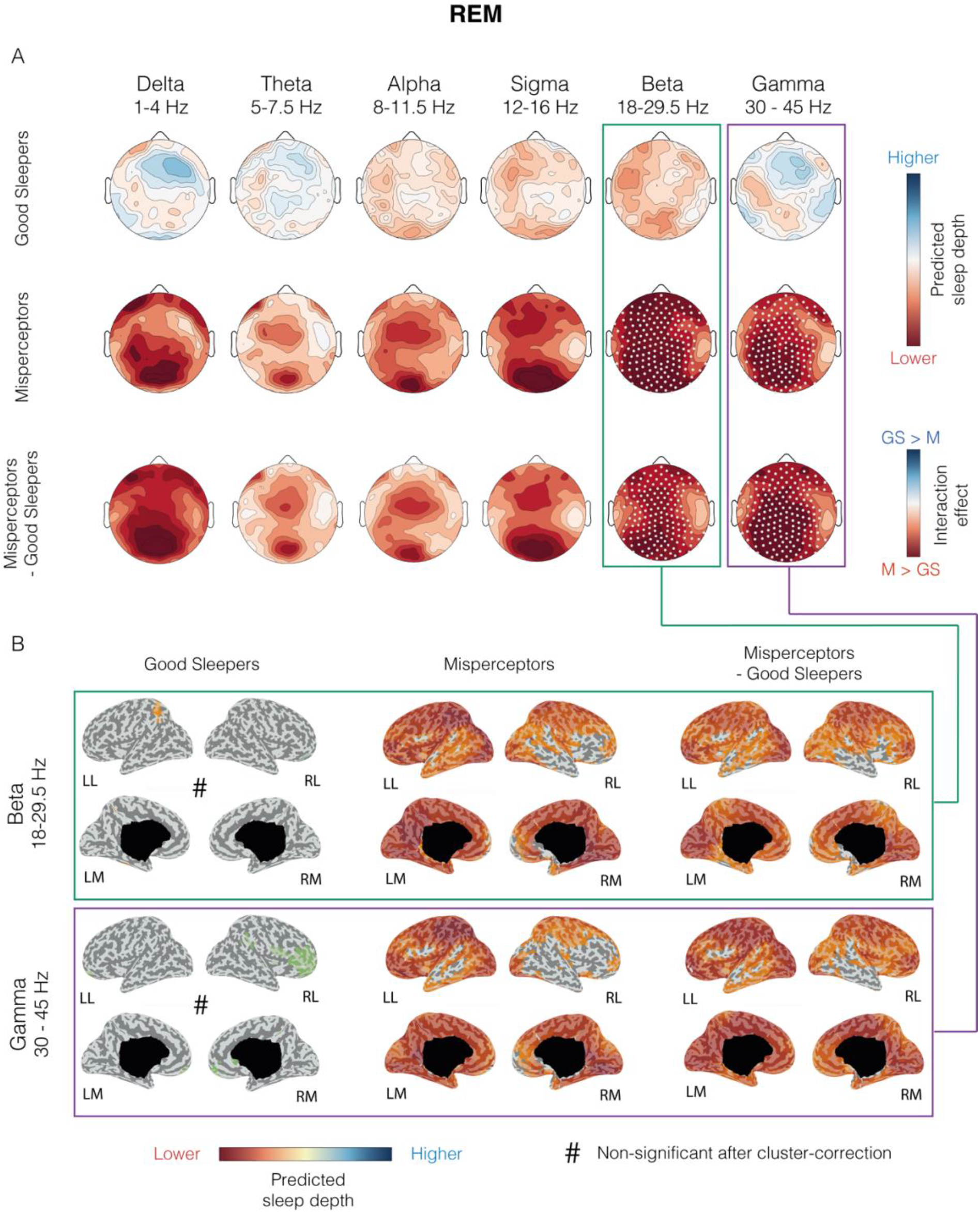
Power spectral density and perceived sleep depth in REM sleep. **A.** Results of linear mixed models explaining perceived sleep depth by power spectral density in good sleepers (n=20, top row) and misperceptors (middle row). Group interaction effect is displayed in bottom row. Power spectral density was averaged over the 120s preceding the awakening. The effect of each frequency band was evaluated in separate models. All models included subject identity and time of night as random factors. Wald statistics values (squared ratio of the fixed factor estimate over its standard error) are shown at the scalp level for each electrode; electrodes with a significant effect after a cluster-and probability-based correction for multiple comparisons (p<0.05) are marked white. **B.** Cortical distribution of Wald statistics at the source level for selected frequency bands (based on results shown in A). The same procedure described in A was used. Voxels with non-significant results are colored in grey. Contrasts that did not survive the correction for multiple comparisons are marked with a hash sign. *LL = left lateral, RL = right lateral, LM = left medial, RM = right medial*.

To understand how spectral power relates to the observation that subjective sleep depth in good sleepers is highest in REM sleep, a sleep state in which the EEG is traditionally described as ‘fast’ and ‘wake-like’, we directly compared spectral power between REM and NREM sleep (**Figure S5**). As expected, low-frequency power (1-16 Hz) was lower in REM sleep compared to NREM sleep. However, high-frequency power in the beta (18-29.5 Hz) and gamma range (30-45 Hz) was not significantly different between the two stages, and in some frontal electrodes even significantly higher in NREM sleep compared to REM sleep. These findings indicate that REM sleep is ‘wake-like’ in terms of low low-frequency power but does not necessarily display higher high-frequency values compared to NREM sleep.

## Discussion

### Sleep perception in good sleepers

Both the feeling of being asleep as well as subjective sleep depth (1 to 5 scale) showed considerable variability within the same night of sleep. Good sleepers reported feeling awake despite full-fledged sleep in ~ 10% of probes, most frequently so during the first two hours of NREM sleep, and again towards the end of the night, during the last hour of sleep. Perceived sleep depth mirrored this course. These observations suggest that the feeling of being asleep sets in progressively, sometime after objective sleep onset, and fades out gradually, before the re-establishment of full wakefulness. These changes in subjective sleep depth are thus not captured by the standard sleep stage classification. The observation that early NREM sleep was associated with the highest proportion of feeling awake instances and among the lowest sleep depth ratings is surprising, since early NREM sleep, displaying the highest amount SWA, is generally considered the “deepest” sleep. This cannot be ascribed to the serial awakening paradigm we used, since in our dataset, SWA decreased progressively across the night (**Figure S.3**), similar to an undisturbed night of sleep, and displayed the highest values during the first two hours of sleep. Even when accounting for the time of the night, we did not find a relation between subjective sleep depth and SWA. These results therefore suggest that SWA and subjective sleep depth can be dissociated. The fact that subjective sleep depth was highest in the middle of the night, near the circadian nadir, could indicate that circadian factors, instead of homeostatic sleep pressure (which is reflected in SWA) may play a role in subjective sleep depth ^29,30^, although our study was not designed to evaluate such circadian effects. Another dissociation between subjective sleep depth and SWA is supported by the observation that REM sleep, in which SWA is reduced and highly localized ^31–33^, featured the highest sleep depth ratings and only 1 out of 126 FAW instances. Since we controlled for the time spent asleep before the awakening, the high subjective sleep depth in this stage cannot be merely a consequence of having spent more time asleep in this stage than in NREM sleep before being awakened.

While we did not find a relation between subjective sleep depth and SWA, we documented a negative relation between high-frequency power in the sigma and beta range and sleep depth in NREM sleep. Consistent with these findings, the course of beta power across the night followed a U-shaped course, inversely mirroring subjective sleep depth (**Figure S.3**). Again consistent with subjective sleep ratings, NREM sleep displayed equal or even higher high-frequency (beta and gamma) power in some brain regions compared to REM sleep (**Figure S.4**). Abrupt increases in high-frequency EEG activity are the defining features of micro-arousals of sleep, which can occur either spontaneously or in response to sensory stimuli, and can be associated with slow waves. It is therefore conceivable that the high-frequency increases that predict lower sleep depth reflect a (subthreshold) activation of arousal systems. The spatially diffuse distribution of sigma and beta power changes are consistent with the widespread projections of arousal systems ^34^. In good sleepers, EEG power did not significantly predict subjective sleep depth in REM sleep, perhaps because of the overall high ratings of subjective sleep depth in this stage and group.

The finding that some spindle parameters predicted lower sleep depth is surprising, since spindles are usually not overtly present in arousals and some rodent studies have shown that periods of increasing spindle activity are associated with increased arousal thresholds^35,36^.

Intra-spindle frequency has been suggested to reflect levels of thalamic polarization within a relatively narrow range ^37^. Spindle frequency typically decreases in the transition to sleep, and fast spindles tend to occur during slow wave up-states during which thalamocortical neurons are depolarized ^38,39^. It is thus possible that relative thalamo-cortical depolarization, closer to the ‘wake state’, which is reflected in fast spindle density, may account for subjectively lighter sleep. Increased spindle amplitude is more difficult to interpret. Although our detection algorithm identified mostly typical spindles, as shown in the average topographical maps (Fig. 4), we cannot exclude that it occasionally picked up high-frequency power increases that were not spindles, but instead arousal-related activity. Indeed, although only sigma and beta levels were significantly related to lower sleep depth (Fig 3.A), similar non-significant results were obtained for a broad range of frequencies from alpha upwards. In addition, centro-parietal electrodes, in which fast spindle density was highest ^38^ displayed the weakest relationship with sleep depth compared to other electrodes.

In line with previous studies, we found that experiences reported upon awakening from REM sleep were more perceptual, richer, longer and associated with lower degrees of lucidity compared to NREM experiences ^26,40,41^. The richer, the more complex and more perceptual the experience even in NREM sleep, the more participants felt asleep. These reports are in line with previous studies showing that thinking and reflective consciousness (comprising lucidity and voluntary control) are typical features of wake mentation, while sleep experiences, especially REM sleep dreams, are highly perceptual ^10,12,26^. In addition, the disappearance of control over thoughts has been shown to relate to the perception of having fallen asleep ^42^. We previously showed that perceptual aspects of dreams correlate with high-frequency power in posterior brain regions, while thought-like experiences correlate with high-frequency power in anterior brain regions ^23^. Here we showed that high-frequency EEG activity predicted lower sleep depth in almost all cortical regions except for parts of the occipital lobe, involved in the perceptual aspects of dreams. It is thus possible that relative frontal “deactivation” ^43^, brought about by quiescence of arousal systems, allows for highly perceptual experiences to emerge, which are associated with feeling deeply asleep.

### Sleep perception in sleep misperceptors

As expected ^16^, sleep misperceptors felt more frequently awake during sleep than good sleepers and perceived their sleep as shallower. The relative distribution of FAW instances between REM and NREM sleep was similar between the two groups, and subjective sleep depth in NREM followed a comparable course across the night (no interaction effects). In addition, in NREM sleep, beta power predicted lower sleep depth like in good sleepers. These findings indicate that feeling awake could be mediated by the same neurophysiological mechanisms in NREM sleep in the two groups, which appear enhanced in sleep misperceptors, possibly as a consequence of hyperarousal (as suggested by metabolic studies in insomnia patients ^44,45^ and animal models of insomnia ^46^). These results are in line with studies that have compared EEG spectral power of whole-night recordings between insomnia patients and good sleepers, and have documented higher relative high-frequency spectral power, comprising the alpha (8-12 Hz), sigma (12-16 Hz)) or beta (16-32 Hz) frequency bands ^25,47–49^, and/or lower relative low-frequency power including the delta and theta band ^49–52^ in insomnia patients. Some studies have extended these findings to insomnia patients with sleep misperception ^4,50,51^. Our results on spindles are also in line with a previous study, showing faster sleep spindles in insomnia patients compared to controls, albeit irrespective of the presence of sleep misperception ^49^.

Unlike good sleepers, who felt most asleep in REM sleep, sleep misperceptors did not display differences in subjective sleep depth between REM and NREM sleep, suggesting that the neurophysiological mechanisms that mediate the feeling of being awake may abnormally persist into REM sleep only in sleep misperceptors. These findings are consistent with a previous serial awakening study showing that patients with insomnia felt more frequently awake than control subjects in REM sleep, but not in N2 sleep ^13^.

Some studies have shown that REM sleep fragmentation by arousals is a prominent feature of insomnia that contributes to the overestimation of wake time during the night ^53^ and interferes with the overnight resolution of emotional distress ^54^. In addition, REM sleep fragmentation in insomnia sufferers has been related to thought-like nocturnal mentation ^54^. In our dataset, insomnia sufferers displayed more thought-like nocturnal mentation compared to controls, irrespective of sleep stage. In addition, 13.8 % of reports of misperceptors contained thoughts and worries about falling alseep, which was never the case for good sleepers. In sleep misperceptors, high-frequency EEG changes predicting sleep depth were more spatially widespread compared to good sleepers, as they involved additional fronto-polar and posterior brain areas, suggesting a more extensive recrutement of arousal systems.

### Overall interpretation

Our results suggest that subjective sleep depth is inversely related to a neurophysiological process that predominates in NREM sleep early in the night, becomes silent in REM sleep, is reflected in high-frequency EEG activity and associated with thought-like conscious experiences. Our results also indicate that this neurophysiological process is amplified in sleep misperceptors: it comes into play more frequently and more strongly (more FAW instances, lower sleep depth), recruits more brain regions (trend towards spatially more diffuse changes) and abnormally persists into REM sleep in this group (no REM-NREM difference in subjective sleep depth). A positive relationship between subjective sleep depth and traditional hallmarks of sleep was not found (sleep spindles, SWA). For sleep spindles, we observed rather an opposite relationship. Thus, it is not the presence of sleep rhythms (slow waves, spindles) but rather the absence of ‘wake-like’ activity (high-frequency power) that appears to account for feeling asleep.

Our data does not provide direct information regarding the origin of the underlying neurophysiological processes, but several of our findings suggest that the noradrenergic arousal system could be a potential mediator of feeling awake during sleep and of low subjective sleep depth. When averaged within behavioral states, noradrenaline levels and locus coeruleus (LC) activity are highest in wakefulness, intermediate in NREM sleep, and lowest during REM sleep, paralleling the incidence of feeling awake instances ^55,56^. Although there are many other neuromodulator systems that show similar profiles across behavioral states (i.e. the histaminergic, orexinergic and serotoninergic systems), several observations make noradrenaline one of the more likely candidates. First, despite overall low LC activity levels in sleep, several studies have shown that the LC discharges intermittently during NREM sleep ^57,58^. Second, while the LC may promote EEG activation and arousal, some studies have documented transient increases in LC activity, reachingquiet wake levels, during full-fledged slow wave sleep at the beginning of the night, when FAW preferentially occurred and subjective sleep depth was lowest in our study ^59–61^. These findings suggest that LC activity can be high even during full-fledged slow wave sleep and thus may account for the dissociation between high SWA and low subjective sleep depth. Another dissociation between SWA and noradrenaline has been documented in fur seals, in which cortical and subcortical noradrenaline levels, unlike EEG activity, do not show lateralization during uni-hemispheric slow wave sleep ^61^. Finally, our source analysis results revealed that high-frequency power changes related to sleep depth were spatially widespread, consistent with the diffuse cortical projections of the noradrenergic system. They tended to spare parts of the temporal and occipital cortices, especially in good sleepers, which is in line with studies showing that the occipital cortex is one of the regions with the lowest levels of cortical noradrenaline ^62,63^, while serotoninergic and some dopaminergic markers display different or opposite cortical distributions^62^. While noradrenergic activity has been linked to the memory benefits of NREM sleep in good sleepers ^64^, its excessive activity has been suggested to disrupt REM sleep in insomnia patients and to lead to dysfunctional emotion regulation ^65^. The inverse relation between subjective sleep depth and spindle amplitude is more difficult to reconcile with noradrenergic activity, as LC firing is known to shorten spindle duration and decrease sigma power, at least in rodents ^66,67^. As discussed above, we currently cannot exclude that our spindle detection algorithm may have mistaken some high-frequency bursts for spindles. Future studies should try to determine whether the reduction of LC activity, for instance with alpha-2 agonists, leads to an increase in subjective sleep depth and how it relates to EEG features in humans ^65^.

### Limitations

Sleep misperceptors represented a relatively small group of highly selected insomnia patients, in which sleep perception was evaluated using a cut-off determined based on the general population. Therefore, the generalizability of findings to other insomnia patients (for instance those with more moderate sleep misperception) is limited. On the other hand, the primary aim of our work was not to understand insomnia, but rather to identify the determinants of sleep perception in general. The extensive probing of experiences through serial awakenings allowed us to study a wide range of sleep perceptions. Including patients with sleep misperception allowed us to extend this range even more and to consider the full spectrum of sleep perception, ranging from ‘normal’ to ‘pathological’. The observation that the determinants of subjective sleep depth in Non-REM sleep are very similar between good sleepers and sleep misperceptors suggests that they are related to sleep perception itself and not insomnia. Our findings in REM sleep on the other hand, reveal interesting differences between the groups that should be further investigated by future studies, which should include insomnia patients with a broader range of sleep misperception.

### Conclusions and perspectives

Our results challenge the commonly held notion that slow wave sleep, generally referred to as deep sleep, best accounts for feeling deeply asleep. Instead, our findings suggest that the subjective perception of sleep is more closely related to the absence of high-frequency EEG activity: the lower it is, the more one feels asleep. Our results also indicate that the feeling of being deeply asleep is associated with perceptual, dream-like experiences, which most often occur in REM sleep, but can also occur in NREM sleep late in the night.

In light of these and other recent findings, one should reconsider the use of the term ‘deep’ sleep to designate slow wave sleep. In fact, NREM sleep is more heterogeneous than previously thought. It comprises alternating phases of sleep stability and fragility and is associated with intermittent noradrenergic surges ^35,57,58,68,69^. Once considered largely dreamless, it is now known to be associated with mental activity in up to 40-70% of cases ^26,28^. Conversely, markers of sleep inertia and high arousal thresholds, traditionally regarded as a hallmark of slow wave sleep, can be seen to a similar extent in REM sleep ^70–72^ and according to some studies^73^, REM sleep appears to better account for subjective sleep quality than slow wave sleep. Not surprisingly, some early sleep electrophysiologists observing cats in different behavioral states chose the term ‘deep sleep’ to refer to REM sleep instead of slow wave sleep ^74,75^. Continuing to use the term ‘deep sleep’ for slow wave sleep could also be misleading for future studies, which may focus exclusively on slow wave enhancement to improve subjective and restorative aspects of sleep, and neglect REM sleep. Finally, the existence of an objective EEG correlate of subjective sleep depth, which is present in both good sleepers and insomnia patients to a different degree, suggests that so-called ‘misperceptors’ are not truly ‘misperceptors’ but may instead accurately perceive subtle shifts towards ‘wake-like’ activity during sleep, which are not apparent in standard sleep recordings.

## Supporting information

All supplementary material

## Acknowledgements

This work was supported by the Swiss National Science Foundation (Ambizione Grant PZ00P3_173955 to F.S.), the Divesa Foundation Switzerland (F.S.) and the Pierre-Mercier Foundation for Science (F.S.). The authors thank, in alphabetical order, David Albir, Françoise Cornette, Stéphanie Dutoit, Grégoire Gex, José Haba-Rubio, Raphaël Heinzer, Eric Lainey, Gianpaolo Lecciso, Nicolas Petitpierre and Tifenn Raffray and the whole CIRS team for patient referral and help with data acquisition; Giulio Bernard, Chiara Cirelli and Giulio Tononi for their comments on earlier versions of the manuscript; as well as Giandomenico Iannetti and Anita Lüthi for their comments on the current version of the manuscript.

## Author contributions

Conceptualization: FS, Methodology: FS (serial awakening paradigm), AS (analysis algorithms). Software and formal analysis: AS and SL. Investigation: JC. Writing – Original draft: FS and AS, Writing – Review & editing: all authors, Visualization: AS, Supervision and project administration: FS, Funding acquisition: FS.

## Star methods

### Selection of participants

Good sleepers (n=20, age 38,3 ± 7,4 yrs (mean±SD), range: 25 – 51, 15 females) were recruited through advertisement and word by mouth. They had to have regular bed and rise times and a good subjective sleep quality (Pittsburgh Sleep Quality Index^76^ <5). Subjects with extreme chronotypes (Horne and Ostberg morningness-eveningness questionnaire^77^ scores >70 or <30), excessive daytime sleepiness (Epworth Sleepiness Scale^78^ >10), suffering from neurological, psychiatric or medical disorders affecting sleep, taking regular medication besides birth control and pregnant females were not included in the study.

Sleep misperceptors of similar age and gender distribution (n=10, age 40,8 ± 6,5 yrs, range: 29 – 49, 8 females, t-test p=0.4) were recruited among outpatients of the Center for Investigation and Research in Sleep at the Lausanne University Hospital. They had to have a diagnosis of chronic insomnia fulfilling the criteria of the international classification of sleep disorders ^79^. Insomnia duration was 15.2 ± 7.8 years (range 2-25). Only patients who had had a polysomnography for clinical reasons were included, which had to fulfil the following criteria: 1) normal sleep efficiency (>85%), 2) sleep latency <60 min. As a routine clinical procedure, immediately after the clinical polysomnography, patients had to estimate how much time they had spent asleep during the night. Only subjects who presented a mismatch between subjective and objective total sleep time (TST) > than 60% were included. The mismatch was evaluated by means of the Sleep Perception Index, which expresses the ratio between subjective (sTST) and objective TST (oTST) in percent ([sTST/oTST] *100). The cutoff of <60% was chosen based on a previous study, in which a sample of the general population (n=2092) underwent polysomnography and estimated TST the next day. In this population, an SPI lower than 58.82% defined the lowest 2.5% of the population distribution. The sample of the general population was comparable, in terms of subjective complaints, PSG and spectral EEG parameters, to the population of insomnia sufferers of this study (for a comparison, see ^4^). Exclusion criteria for patients comprised major psychiatric and neurological comorbidities, medication other than birth control, pregnancy, apnea-hypopnea index >15/h and periodic leg movement index in sleep >15/h. Written informed consent was obtained by all the participants and the study was approved by the local ethical committee (commission cantonale éthique de la recherche sur l’être humain du canton de Vaud).

As part of the work-up, all participants completed the Beck Anxiety and Depression Inventories (BAI and BDI respectively). Average values for the two populations were within the norm (<7 points for the BAI and <9 points for the BDI). All subjects scored below or within limits of mild depression or anxiety (8-15 for BAI, 10-18 for BDI) except for two sleep misperceptors, who had moderately elevated scores. Patients with insomnia had significantly higher depression scores than good sleepers (6.1±4.6 vs 1.4±2.3 points, p=0.011, two-tailed paired t-test) but comparable anxiety scores (2.8±3.8 vs 7.3±7.5 points, p=0.011). Neither BDI nor BAI scores correlated with the probability of feeling awake (FAW) [Spearman correlation BDI: rho=0,35, p=0,13; BAI: rho=0,29, p=0,22] or SPI [BDI: rho=-0,26, p=0,27; BAI: rho=-0,05, p=0,84].

### Experimental procedure

Both sleep misperceptors and good sleepers underwent an undisturbed baseline hd-EEG sleep recording with 256 electrodes in the sleep laboratory, starting at approximately 11:30 pm and ending at 6:30 am the next morning. Sleep architecture and EEG findings of this baseline night (same 10 sleep perceptors and 24 controls) have been published separately ^4^. In brief, sleep parameters in insomnia sufferers did not differ significantly from controls, except for a significantly lower proportion of stage N3 sleep ^4^. Participants subsequently underwent two experimental hd-sleep EEG recordings, in which a a previously validated serial awakening paradigm was employed ^26^. The two experimental nights were scheduled at least one week apart to limit the effects of the experimental sleep fragmentation on dream recall. One good sleeper completed only one experimental night because of organizational constraints. Subjects were asked to maintain a regular sleep-wake schedule during the week before the recordings, and compliance was verified with wrist-worn actigraphy. Participants did not have access to watches or phones during the recordings.

Awakenings were carried out with a 1.5s computerized alarm sound that was played through intercom by the examiner who continuously monitored the subjects via EEG and video/audio from a control room ^26^. NREM awakenings were performed approximately 10 min after the first sleep spindle, while rapid eye movement (REM) sleep awakenings were carried out 10 min after the beginning of REM sleep.

After each awakening, participants were asked to estimate whether they had been feeling awake or asleep before the alarm sound. Next, they had to estimate, on a 1-5 scale, how deeply they had felt asleep before before the alarm sound (depending on the answer to the previous question). In all cases, they also had to describe what they had in mind immediately before the alarm sound (“What was the last thing going through your mind prior to the alarm sound?”). The presence of a conscious experience (CE) was considered separately from instances in which subjects reported conscious experiences but could not report any content (CE without report of content, CEWR), and from no experiences (NE). When an experience was reported, participants had to estimate the following dimensions on a 1 to 5 point scale: 1) the degree of lucidity (“How much did you realize that the experience was happening in your mind and not in reality?”), 2) degree of voluntary control over the experience (“How much did you have control over the content of the experience?”), 3) the thought-like character of the experience (“How much did it resemble a thought or reflection ?”), 4) the perceptual character of the experience (“How much was it perceptual?”), 5) the length of the experience (“How long was the experience ?”) as well as 6) the richness and complexity of the experience (“How rich and complex was the experience ?”). Once the interview was finished, subjects were allowed to fall back to sleep. No feedback was given on whether they had been awake or asleep. Reports were reviewed retrospectively by an experimenter to quantify the presence of sleep-related thoughts and preoccupations.

### Sleep scoring and EEG preprocessing

A 256-channel system (Electrical Geodesics, Inc., Eugene, Oregon) was used for EEG acquisition, with a sampling rate of 500 Hz. Four of the 256 electrodes near the eyes were used to monitor eye movements, electrodes overlying the masseter muscles and close to the chin were used to monitor muscle tone ^23^. The EEG signal was off-line band-pass filtered between 0.5 and 45 Hz, and sleep scoring was performed over 30s epochs according to standard criteria ^80^.

The two minutes of data before each awakening were extracted for analysis. Channels containing artefactual activity were visually identified and replaced by interpolation using information from the remaining channels using spherical splines (NetStation, Electrical Geodesic Inc.). Ocular, muscular, and cardiac artefacts were eliminated with Independent Component Analysis (ICA) using EEGLAB routines ^81^. ICA components with specific activity patterns and component maps characteristic of artefactual activity were removed ^82^.

To determine the time spent asleep before each awakening, all the recordings were visually inspected. Wake periods were scored as at least 15s of clear-cut EEG wakefulness ^80^. The time between the end of the last period scored as wake and the alarm sound was extracted for each awakening (= time spent asleep). All subsequent EEG analyses were performed on average-referenced data.

### EEG analysis

Power spectral densities (PSD) were calculated using the Fast Fourier transform method on artifact-free consecutive, non-overlapping 2s epochs and used to compute signal power in typical frequency bands, including delta (1–4 Hz), theta (5–7.5 Hz), alpha (8–11.5 Hz), sigma (12–16 Hz), beta (18–29.5 Hz) and gamma (30 Hz – 45 Hz). In order to avoid overlap between frequency bands, we excluded the upper 0.5 Hz range of neighbouring bands.

Source localization was performed using a previously described procedure ^23^. In brief, the cleaned and filtered signal corresponding to the 2min of sleep period prior to each awakening were extracted. Source modeling was performed using the GeoSource software (Electrical Geodesics, Inc., Eugene, Oregon). Individualized geocoordinates of the electrodes were used to construct the forward model. The source space was restricted to 2,447 dipoles that were distributed over 7×7×7-mm cortical voxels. The inverse matrix was computed using the standardized low resolution brain electromagnetic tomography (sLORETA) constraint ^83^. A Tikhonov regularization (λ=10-2) procedure was applied to account for the variability in the signal-to noise ratio ^83^.

Spindle detection was based on an automatic algorithm, adapted from a previous study ^84^. The EEG signal was average-referenced, downsampled to 128 Hz and bandpass-filtered between 11 and 16 Hz (~3 dB at 10 and 17 Hz). The following spindle parameters were extracted: density, density of low and high frequency, maximal amplitude, duration and frequency. Based on previous studies ^37,38,85^, we categorized spindles as fast or slow based on an individualized threshold, which was defined as the intermediate value between the average spindle frequency in one centroparietal channel (Pz) and one frontal channel (Fz) ^86^.

### Statistical analysis

Statistical analyses were performed in R-Studio (version 1.1.463). In the following section and in the result section, reference numbers preceded by hash signs refer to the statistical models listed in Table 1.

Generalized mixed models were computed to evaluate the predictors of perceived state (FAW vs FAS, categorical dependent variable), including group and stage (#1, Fig.1A), group and time of night (hours after lights off) in NREM (#2, Fig.1B), and group and conscious experience presence and recall (#6, Fig.2A). Linear mixed models were computed to evaluate the predictors of perceived sleep depth (Degree, ordinal dependent variable), including group, stage and time (#3, Fig.1C and Fig.1D), group, stage and C.Exp (#8) and group, stage and dream content characteristics (#9). Linear mixed models were also computed to evaluate the predictors of dream content characteristics (listed in the experimental procedure section), including group and sleep stage in FAS (#5, Fig.2B) and group and perceiveed state in NREM (#7, Fig.2D). A generalized mixed model was computed to evaluate the effect of group and stage of night on presence or absence of conscious experience (CE_presence, categorical dependent variable, #4, Fig.2C). Subject identity and time spent asleep before each awakening were included as a random factor in all analyses. Post-hoc contrasts were computed with estimated marginal means (emmeans package) and a Tukey’s HSD (honestly significant difference) test was applied to adjust for the risk of false discoveries (type I error) consequent to multiple comparisons. In the results section, all main statistics and associated p-values are reported in the text, post-hoc statistics are displayed in the figures when possible, and all estimates and post-hoc values are reported in supplementary tables referenced in the text.

Pearson correlation was performed in order to assess the correlation between the proportion of FAW in the two serial awakening nights and between the proportion of FAW in the serial awakening night and the sleep perception index in the baseline night. In order to control for the difference in range of proportion of FAW and SPI between groups, these correlation analyses were performed on data z-scored within groups.

For EEG data analyses, linear mixed models were computed for each electrode (at the scalp level) or voxel (at the source level) to evaluate the effect of absolute power spectral density in different frequency bands (#10, Fig.3 and 4) and different spindle parameters (#11, Fig.6) on perceived sleep depth. For models #10 and #11, for each frequency band or spindle parameter, a separate model was computed. Subject identity and time of night was included as a random factor in all the analyses, and frequency bands and group as fixed factors. As mixed model results display the effect of each fixed factor considering all other fixed factors at reference value, we performed the models twice with each group as reference in order to obtain scalp topographies of model effects for each group. As between-subject variability is accounted for by including subject identity as random factor, all EEG power spectral density models were performed on absolute (non-normalized) power values. For each model, we obtained the Wald statistics, which is the squared ratio of the fixed factor estimate over its standard error. To account for multiple comparisons, a cluster and probability-based correction was used. In order to create a dummy population, labels corresponding to the dependent variables were shuffled within subjects 1000 times. For each permutation, the model was applied and neighboring electrodes with p-values inferior to 0.05 were identified as a cluster. A cluster statistics was produced by summing Wald statistics inside each significant cluster and by dividing by the number of electrodes in the cluster, resulting in one value for each identified cluster. For each permutation, only the absolute maximal value of all cluster statistics was retained. A threshold for significance was set at the 95th percentile of the dummy cluster statistics distribution. The cluster statistics obtained in the real dataset were considered significant when above this threshold. This procedure was applied for each electrode/voxel.

