## Supplementary material for "The determinants of subjective sleep depth: insights from a high-density-EEG study with serial awakenings": All supplementary material

### SUPPLEMENTARY FIGURES

A

**Quantity of trials per group, stage and perceived state**

| | GS | | | SM | | |
|---|---|---|---|---|---|---|
| | **FAW** | **FAS** | **TOTAL** | **FAW** | **FAS** | **TOTAL** |
| **NREM** | 50 | 358 | 408 | 76 | 116 | 192 |
| **REM** | 1 | 125 | 126 | 6 | 55 | 61 |

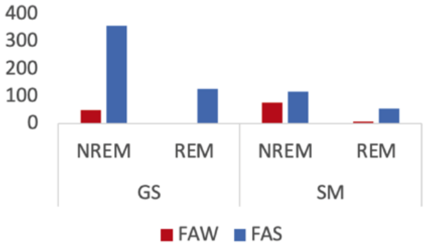

B

**Percentage of FAW/FAS per group and stage**

| | NREM | | REM | |
|---|---|---|---|---|
| | **FAW** | **FAS** | **FAW** | **FAS** |
| **GS** | 12,3 | 87,7 | 0,8 | 99,2 |
| **SM** | 39,6 | 60,4 | 9,8 | 90,2 |

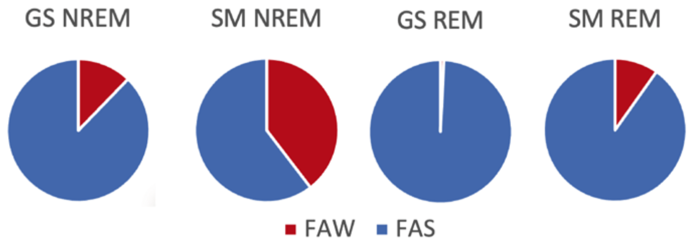

C

**Percentage of conscious experiences per group and stage**

| | GS | | SM | |
|---|---|---|---|---|
| | **NREM** | **REM** | **NREM** | **REM** |
| **NE** | 21,3 | 10,3 | 25,0 | 13,1 |
| **CEWR** | 38,7 | 26,2 | 41,7 | 27,9 |
| **CE** | 40,0 | 63,5 | 33,3 | 59,0 |

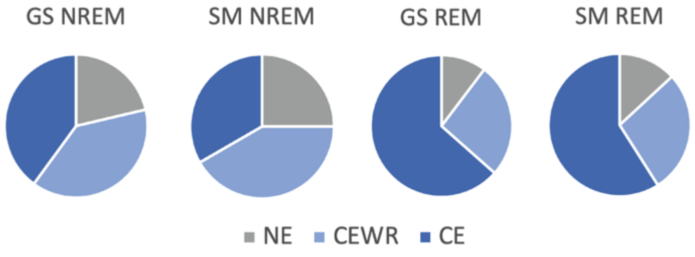

**Figure S.1. Distribution of awakenings (trials) across group, stage, perceived state and conscious experiences. A.** Table and bar graph of quantity of awakenings per group, stage and perceived state. **B.** Table and pie chart of percentage of FAW and FAS per group. **C.** Table and pie chart of percentage of conscious experiences per group and stage. *CE = Conscious experience with recall; CEWR = Conscious experience without recall; FAS = Feeling asleep; FAW =* Feeling awake; *GS = Good sleeper; NE = No experience; NREM = Non-REM sleep; REM = Rapid-eye movement sleep; SM = Sleep misperceptors;*

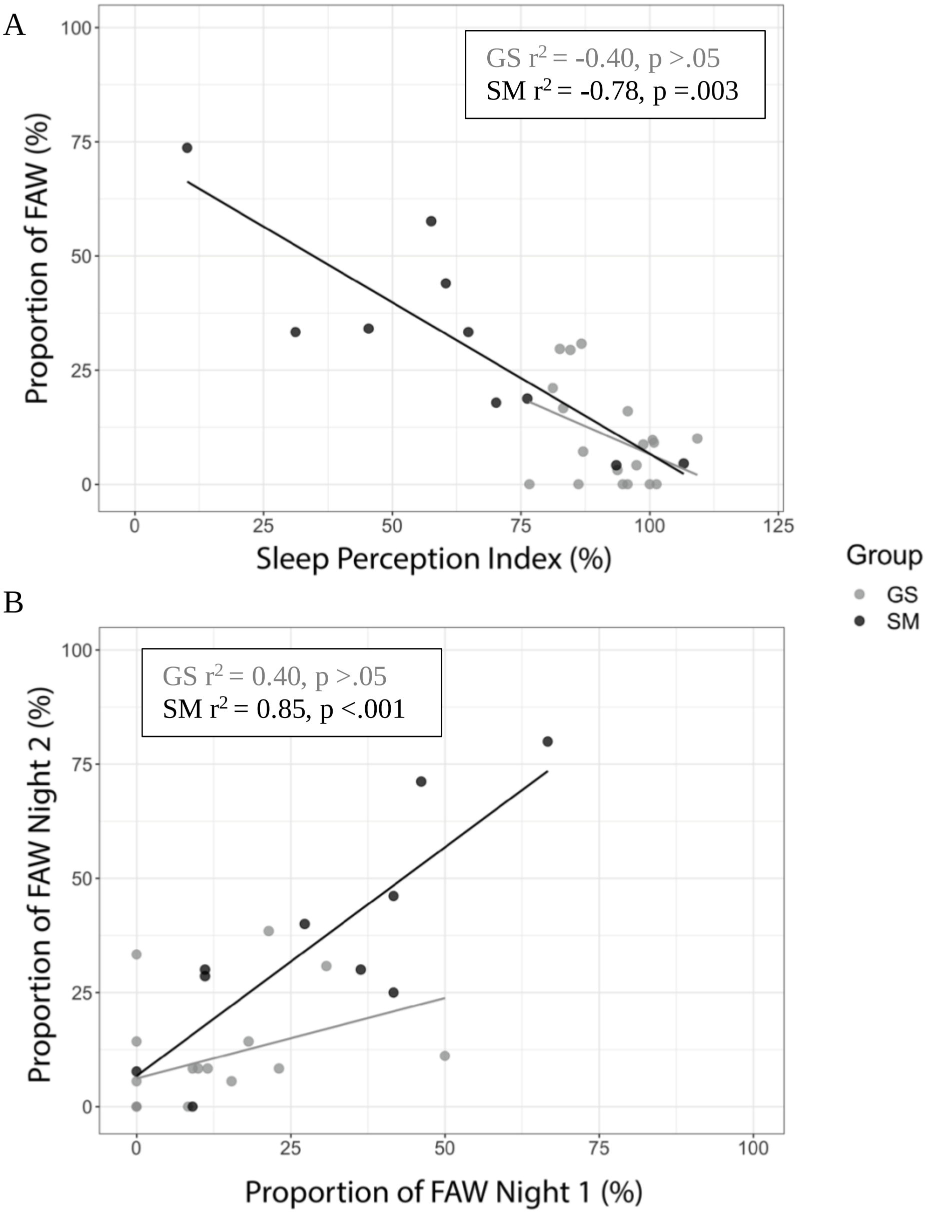

**Figure S.2. Correlation between sleep misperception measures in serial awakening and baseline nights. A.** Pearson correlation between the average proportion of FAW in the two serial awakening nights and the sleep perception index during an uninterrupted baseline night in good sleepers (GS, n=20; grey) and sleep misperceptors (SM, n=10, black). One trend line per group is displayed on the figure, as well as the $r^2$ and corresponding p-value. **B.** Correlation between the average proportion of FAW in the two serial awakening nights in good sleepers (GS, n=19; grey) and sleep misperceptors (SM, n=10, black). For both analyses, Pearson correlation was performed on data that was z-scored within group to control for the difference of range of SPI and FAW proportion between groups. To ease interpretation, original non z-scored values are displayed in the plots.

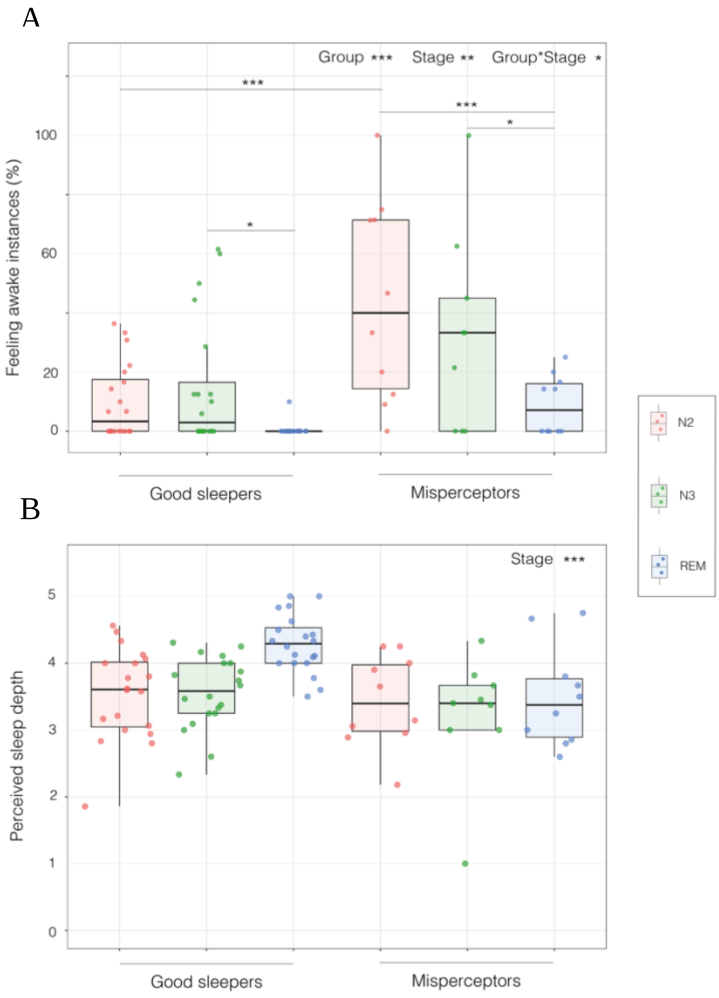

**Figure S.3. Determinants of sleep perception in N2, N3 and REM sleep.** Perceived state (feeling awake vs feeling asleep; A) and degree of perceived sleep depth (scale from 1 to 5; B) as a function of group (good sleepers and sleep misperceptors) and stage (N2, N3, REM sleep). **A.** Proportion of feeling awake instances in good sleepers (n=20, left) and misperceptors (n=10, right) in sleep stages N2 (red), N3 (green) and REM sleep (blue). **B.** Degree of perceived sleep depth in good sleepers (grey) and sleep misperceptors (black) in NREM and REM sleep. Each point represents an individual, the box displays the 25$^{th}$ to 75$^{th}$ percentile of data and the vertical bars shows the confidential interval. Post-hoc of significant main and interaction effects evaluated with generalized linear mixed models are indicated with lines and asterisks indicating the level of significance: * *$p < 0.05$*, ** *$p < 0.01$*, *** *$p < 0.001$*. All main and post-hoc effects are reported in tables 3 and [illegible]

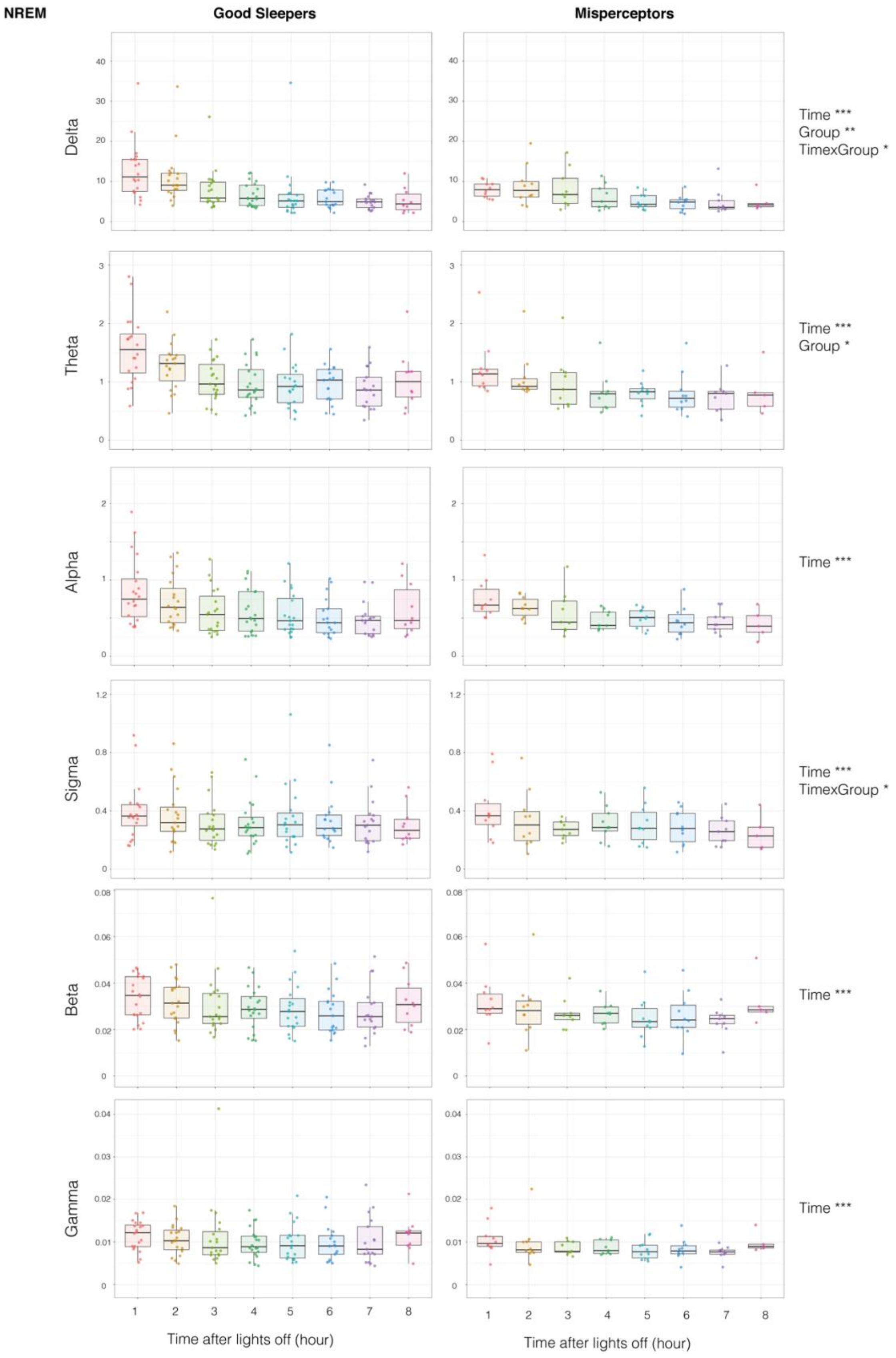

**Figure S.4. Evolution of power spectral density in the course of the night in NREM sleep of serial awakening.** Power spectral density averaged over all electrodes for diverse frequency bands from lowest (Delta; upper line) to highest (Gamma; bottom line) on the two minutes preceding awakening. Each point represents an individual's average power in the course of the night, the box displays the 25th to 75th percentile of data and the vertical bars shows the confidential interval. Main simple and interactions effects of linear mixed models power spectral density by time after lights off – controlling for subject identity and time spent asleep – are indicated on the right of each frequency band plot. * $p < 0.05$, ** $p < 0.01$, *** $p < 0.001$. All main and post-hoc effects are reported in tables 22 to 27.

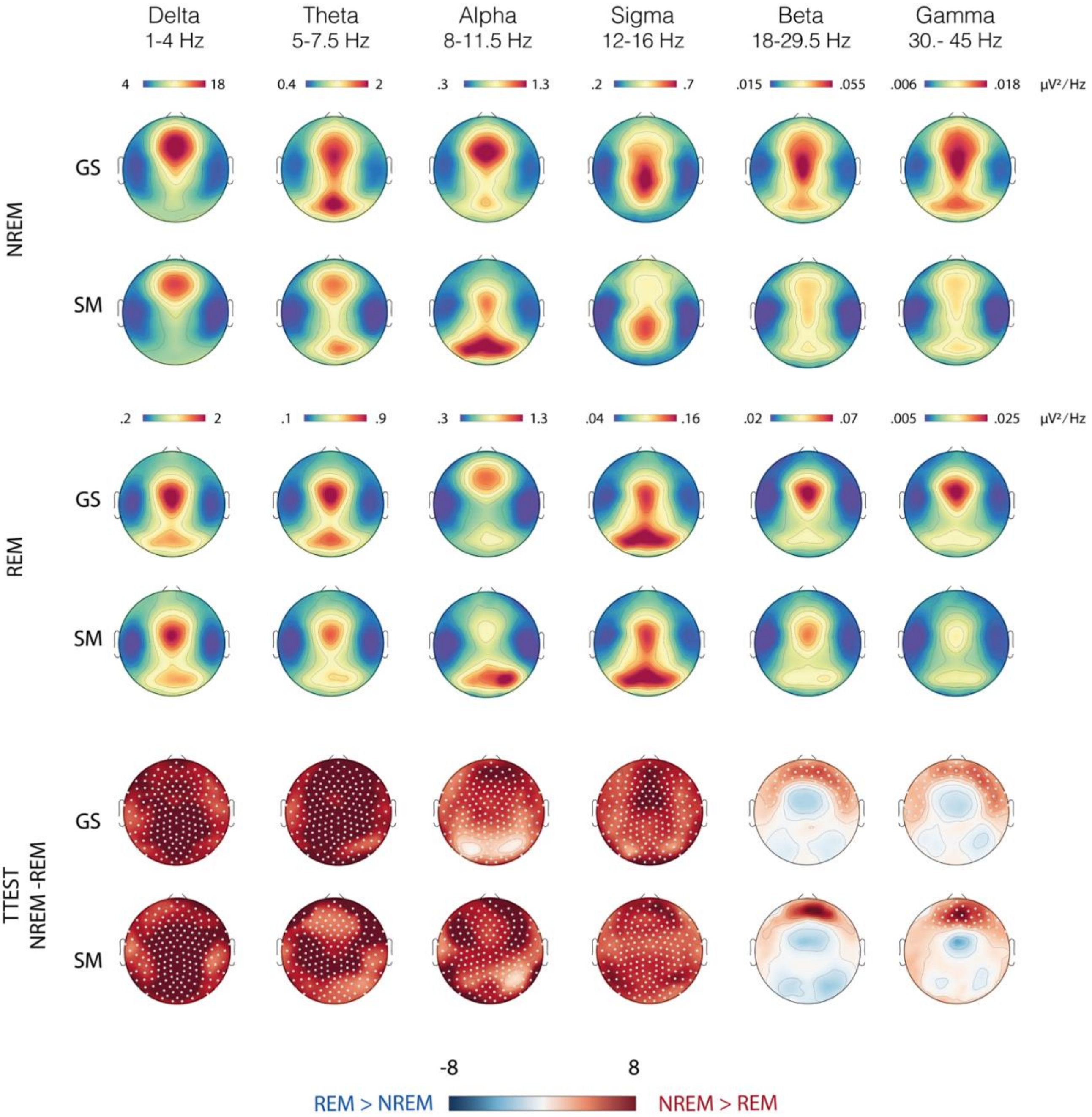

**Figure S.5. Power spectral density in NREM and REM sleep.** Absolute power spectral density of diverse frequency bands (columns) in good sleepers (GS, n=20) and subjects with sleep misperception (SM, n=10) in NREM sleep (top row), REM sleep (middle row). T-values resulting from T-test between REM and NREM sleep power spectral density are displayed in the bottom row. White dots indicate channels with a significant effect of stage after cluster-correction. Cluster correction was performed as described in the method section.

**Annex S.1. Examples of reported experiences associated with feeling awake (FAW) and feeling asleep (FAS)**

The reports below represent responses to the question asked by the examiner after awakening the subject with an alarm: “What was going through your mind just prior to the alarm?”. All FAW reports are shown, as well as a selection of FAS reports (for two representative good sleepers and misperceptors). FAS tables also display the “Degree” of perceived sleep depth (1 to 5 = deepest).

**FAW reports in good sleepers**

| ID | Night | Stage | Time | Report |
|---|---|---|---|---|
| GS1 | 1 | N2 | 00:23 am | I was thinking about something, familiar faces, there were familiar faces. But I cannot remember exactly which ones. |
| GS2 | 1 | N3 | 23:25 pm | I was thinking about my day today. |
| GS2 | 2 | N3 | 22:52 pm | I was thinking about my day today. |
| GS3 | 2 | REM | 06:00 am | I was thinking about patients who cannot comply with their medical treatments, patients who are in unfavorable conditions, who do not have money, who cannot come to the hospital and be treated. |
| GS4 | 2 | N2 | 23:26 pm | I was thinking about my tomorrow. About how to organize my day. I was simply thinking about how the day will unfold, me arriving at the hospital ward, the breaks, and what will happen in the evening. To be very very precise, I see myself entering the hospital and greeting the people there, that’s where I was at, I think. |
| GS5 | 1 | N2 | 00:15 am | I was thinking about the signals related to being awake and being asleep |
| GS5 | 1 | N2 | 00:34 am | Images of a swimming pool. |
| GS6 | 1 | N2 | 03:34 am | I was thinking about the organization of my day. |
| GS6 | 1 | N2 | 04:32 am | I was thinking about an apartment. |

**FAW reports in misperceptors**

| ID | Night | Stage | Time | Report |
|---|---|---|---|---|
| SM1 | 1 | N2 | 00:47 am | I believe I was thinking about what I should do to fall asleep, to find sleep, to relax. |
| SM1 | 1 | N2 | 02:06 am | Mostly thoughts about falling asleep, me falling asleep. Thoughts about the fact that I don't fall asleep. |
| SM1 | 2 | N2 | 00:29 am | I can't wait to fall asleep. |

| | | | | |
|---|---|---|---|---|
| SM2 | 1 | N2 | 23:34 pm | I was thinking about the organization of a project that I have to do. |
| SM2 | 1 | N2 | 00:34 am | I was still thinking about the organization, but I could not tell you more. |
| SM2 | 2 | N2 | 23:48 pm | About organizing something. |
| SM2 | 2 | N2 | 06:15 am | I was saying to myself that I was awake. |
| SM3 | 1 | REM | 02:30 am | That something was preventing me from sleeping, that I felt like stripping off the EEG net. |
| SM3 | 2 | N3 | 01:43 am | Some banalities. Something on the table. In short, I was thinking about how were going to set the table for when there are people. |
| SM4 | 1 | N2 | 00:12 am | I was thinking that I forgot to wear the watch. |
| SM4 | 1 | N2 | 00:44 am | I was telling myself that I would not be able to sleep. |
| SM4 | 2 | N3 | 23:46 pm | I was thinking about things that I have to do, and about what I have to pack for work tomorrow. |
| SM4 | 2 | N2 | 00:18 am | I was thinking about some jobs that I am doing at the moment. |
| SM5 | 1 | REM | 05:56 am | That I was not able to sleep. |
| SM5 | 2 | N3 | 00:21 am | That I want to sleep. |
| SM6 | 2 | N2 | 00:58 am | I was thinking about the study. |
| SM7 | 1 | N3 | 23:57 pm | I was thinking about the house, my son and all that, and whether he was already in bed. |
| SM7 | 1 | REM | 01:07 am | I was thinking about the things that are happening at home, in Portugal. |
| SM7 | 1 | N3 | 02:38 am | I haven’t slept. I am thinking about many things, like little movies in my head, about the electrodes that prevent me from sleeping, it is a little painful, I can’t fall asleep. There are people getting in and out of the metro, people passing. These are not thoughts but images. |

**Representative FAS reports in good sleepers**

| ID | Stage | Time | Degree | Report |
|---|---|---|---|---|
| GS14 | N3 | 11:27 pm | 3 | I don’t remember exactly. Yes I do. I was taking a carpentry class. It was related to blocks of wood interlocking. |
| GS14 | N3 | 0:25 am | 3 | I was cooking, I was taking cooking classes, pastry even. I was explaining to someone how to make pastries. |
| GS14 | N2 | 1:18 am | 4 | The fact that organic vegetables are wrapped in plastic. |

| | | | | |
|---|---|---|---|---|
| GS14 | N2 | 1 :32 am | 3 | I have the feeling that it was closely related to the moment we spoke before. It is a bit vague, but I think I was explaining that I was sleeping as part of a sleep study. |
| GS14 | REM | 2 :45 am | 4 | I was on the phone with my mother, and we were talking about family questions and mourning and that kind of things. |
| GS14 | N2 | 3 :03 am | 4 | There was somebody else with me, not people I know. I can't remember why. There was somebody mocking somebody else actually. |
| GS14 | N2 | 3:17 am | 5 | It was a bit like a movie. It looked a bit like the Big Lebowski and reality. I was a witness of that, I wasn't part of the scene actually. |
| GS14 | REM | 4:28 am | 5 | I was dreaming that there was some sort of mushroom. I was examining the mushrooms that I had picked and then I found myself with my boyfriend and a friend at some sort of village fair, but it was in Scotland and it was a Swiss celebration, so there were stands for all Swiss cantons. We went there and then I woke up. |
| GS14 | N3 | 5:29 am | 5 | It is going to sound weird, but I was only seeing colors actually. Red and purple. |
| GS14 | N3 | 1:15 am | 5 | My little brother who was with a friend of his and I don't remember why now, but he was taking the train, he was traveling by train. |
| GS14 | REM | 5:58 am | 5 | I was getting my hair dyed. |
| GS19 | N3 | 0:12 am | 3 | I was thinking of a green color. |
| GS19 | N3 | 0:33 am | 4 | The shape of noise, determining if it was full or empty, full of hollow. I don't know how to describe it. Whether the lid was full or empty. I dreamt about a lid made of some material and we couldn't tell. |
| GS19 | N3 | 1:08 am | 4 | I would say « tidying up». How to tidy up. |
| GS19 | REM | 2:20 am | 5 | I was seeing the stitches of a sweater, a knit. |
| GS19 | REM | 4:01 am | 5 | I was in a car with my ex-boyfriend, he was driving, I was a bit upset because I had lost my belongings in a fire in my apartment. In the dream, he was actually giving me advice. |
| GS19 | REM | 5:33 am | 5 | I was in a shop with a dog I know, it's a friend's dog, he was fixed to a support, attached actually, he was being mistreated, it was weird. As if it was a model exposed in the shop. So, I got a bit angry, I complained to the shop keeper "What are you doing? This is inacceptable!" and so on. Then I freed the dog and you contacted me. He had velcro straps around his legs and the shop had attached a price tag on one of his paws, which was hard to remove, all his fur came off, it was horrifying. His right eye was in a bad shape, he must have gotten into a fight and gotten hurt. |
| GS19 | N2 | 6:27 am | 5 | There was an alarm clock exhibited, with a glass of water. I drank the glass of water. I cleared the alarm clock because there were brambles all around it. |

**Representative FAS reports in misperceptors**

| ID | Stage | Time | Degree | Report |
|---|---|---|---|---|
| SM3 | REM | 2:30 am | 2 | Something was preventing me from sleeping and I wanted to tear off the net. |
| SM3 | REM | 4:32 am | 3 | I was thinking about some things. A dream. It was some sort of game, a game that we made people play. Like in “The game”, the movie “The game”. |
| SM3 | N2 | 5:22 am | 1 | I was wondering about my sleep. I was thinking about my sleep. |
| SM3 | REM | 6:00 am | 2 | Memories. Some memories. I don’t remember clearly. There were people dressed up, I can’t remember. |
| SM3 | N2 | 6:27 am | 2 | I was thinking again about my sleep. On how to manage. I can’t remember exactly how but it was related to managing my sleep. |
| SM10 | N3 | 0:10 am | 5 | I was dreaming that I was on holiday. That I was planning my holidays. |
| SM10 | N3 | 0:25 am | 5 | I was planning my holidays again. |
| SM10 | N3 | 0:46 am | 5 | I was planning my holidays. |
| SM10 | N2 | 2 :26 am | 4 | I was with my daughter, and she was riding a horse. |
| SM10 | N2 | 2 :45 am | 5 | I had to get up because it was almost time to go to work. |
| SM10 | REM | 3 :46 am | 5 | I was dreaming. In the dream there were two people singing, and I thought it was nice. |
| SM10 | REM | 5 :48 am | 5 | I was dreaming that I was with my husband and my children at home, and that our neighbors were there. They were looking for a traveling agency to plan a week-end trip to Paris. |

#### STATISTICS TABLES

*For each model, the first table present analysis of variance of linear mixed model results and the second table presents model estimates of the model. Analysis of variance was of type II in absence of a significant interaction effect and type III otherwise. In the estimates tables, t values are given for linear mixed models and z values for generalized mixed model. In case of significant main effect with more than two level or a significant interaction effect, an additional table presents the lsmeans pairwise meaningful contrasts performing a Tukey's HSD (Honest Significant Difference) test, adjusting for multiple comparisons. The annotation X1*X2 always means X1 + X2 + X1:X2 where X1:X2 is the interaction effect. The annotation X1*X2*X3 includes all simple effects, all two variable interaction effects and the triple interaction. The "time asleep" factor is calculated as the time between the last wake page and the awakening. For all models, table A displays main effects, table B indicates model estimates, table C shows post-hocs of significant simple terms effect and table D post-hocs of significant interaction terms effect. The number following a hash sign indicates the model ID.*

**Summary**

**TABLE 15 (#6): PERCEIVED SLEEP DEPTH ~ GROUP*STAGE*CE + (1|SUB) + (1|TIME ASLEEP) IN FAS** ... 19
**TABLE 16 (#7A): PERCEIVED SLEEP DEPTH ~ GROUP*STAGE*THOUGHT-LIKE + (1|SUB) + (1|TIME ASLEEP) IN FAS** ........ 20

TABLE 17 (#7B): PERCEIVED SLEEP DEPTH ~ GROUP*STAGE*PERCEPTUAL + (1|SUB) + (1|TIME ASLEEP) IN FAS ........................................................................ 20
TABLE 18 (#7C): PERCEIVED SLEEP DEPTH ~ GROUP*STAGE*RICHNESS&COMPLEXITY + (1|SUB) + (1|TIME ASLEEP) IN FAS ........................................................................ 21
TABLE 19 (#7D): PERCEIVED SLEEP DEPTH ~ GROUP*STAGE*DURATION + (1|SUB) + (1|TIME ASLEEP) IN FAS ........................................................................ 21
TABLE 20 (#7E): PERCEIVED SLEEP DEPTH ~ GROUP*STAGE*CONTROL + (1|SUB) + (1|TIME ASLEEP) IN FAS ........................................................................ 22
TABLE 21 (#7F): PERCEIVED SLEEP DEPTH ~ GROUP*STAGE*LUCIDITY + (1|SUB) + (1|TIME ASLEEP) IN FAS ........................................................................ 22
TABLE 22 (#9A): DELTA ~ TIME + GROUP + TIME:GROUP + (1|SUB) + (1|TIME ASLEEP) IN NREM – FIGURE S.2........................................................................ 23
TABLE 23 (#9B): THETA ~ TIME + GROUP + TIME:GROUP + (1|SUB) + (1|TIME ASLEEP) IN NREM – FIGURE S.2........................................................................ 24
TABLE 24 (#9C): ALPHA ~ TIME + GROUP + TIME:GROUP + (1|SUB) + (1|TIME ASLEEP) IN NREM – FIGURE S.2........................................................................ 24
TABLE 25 (#9D): SIGMA ~ TIME + GROUP + TIME:GROUP + (1|SUB) + (1|TIME ASLEEP) IN NREM – FIGURE S.2........................................................................ 25
TABLE 26 (#9E): BETA ~ TIME + GROUP + TIME:GROUP + (1|SUB) + (1|TIME ASLEEP) IN NREM – FIGURE S.2........................................................................ 26
TABLE 27 (#9F): GAMMA ~ TIME + GROUP + TIME:GROUP + (1|SUB) + (1|TIME ASLEEP) IN NREM – FIGURE S.2........................................................................ 26

**Abbreviations**

| Abbreviation | Meaning |
|---|---|
| C. Exp | Conscious Experience (Categorical Variable) |
| CE | Conscious Experience With Recall of content (C.Exp Level) |
| Chisq | Chi-square statistic |
| Df | Degree of freedom |
| FAS | Feeling asleep (Perceived State Level) |
| FAW | Feeling awake (Perceived State Level) |
| GS | Good Sleepers (Group level) |
| Inf | Standard output of R function emmeans indicating estimates are tested against a standard normal distribution (z-test) |
| IV | Independent Variable |
| NE | No Experience (C.Exp Level) |
| NREM | Non Rapid-Eye Movement Sleep (Stage Level) |
| REM | Rapid-Eye Movement Sleep (Stage Level) |
| SM | Sleep Misperceptors (Group level) |
| Stage | Categorical variable: NREM vs REM |
| Stage_all | Categorical variable: N2 vs N3 vs REM |
| Std Error | Standard error |
| Sub | Subject identity |
| Time | Time in hours after lights off |
| Time_asleep | Time in minutes after last wake |
| Time_since_first_sleep_onset | Time since first N1 |

**I. Effect of group, sleep stage and time of the night on perceived state (FAW/FAS) and perceived sleep depth**

**Table 1 (#1): Perceived state ~ Group*Stage + (1|sub) + (1|time asleep) – Fig.1.A**

**Table 1.A. Main effects**

| IV | Chisq | Df | P value |
|---|---|---|---|
| **Group** | **11,003** | **1** | **0,001** |
| **Stage** | **7,727** | **1** | **0,005** |
| Group:Stage | 0,284 | 1 | 0,594 |

**Table 1.B. Estimates**

| IV | Estimate | Std Error | Z value | P value |
|---|---|---|---|---|
| **(Intercept)** | **-2,263** | **0,334** | **-6,769** | **<0,001** |
| **SM-GS** | **1,786** | **0,538** | **3,317** | **0,001** |
| **REM-NREM** | **-2,838** | **1,021** | **-2,780** | **0,005** |

**Table 1.C. Post-hocs: Simple terms**

| IV | | Estimate | SE | Df | z.ratio | p.value |
|---|---|---|---|---|---|---|
| **Group** | **SM-GS** | **1,64** | **0,50** | **Inf** | **3,31** | **<0,001** |
| **Stage** | **REM-NREM** | **2,37** | **0,44** | **Inf** | **5,35** | **<0,001** |

**Table 2 (#2): Perceived State ~ Group*Time + (1|sub) + (1|time asleep) in NREM sleep – Fig.1.B**

**Table 2.A. Main effects**

| | Chisq | Df | P value |
|---|---|---|---|
| **Group** | **9,026** | **1** | **0,003** |
| **Time** | **30,004** | **7** | **<0,001** |
| Group:Time | 7,786 | 7 | 0,352 |

**Table 2.B. Estimates**

| | Estimate | Std Error | z value | P value |
|---|---|---|---|---|
| **(Intercept)** | **-1,263** | **0,515** | **-2,453** | **0,014** |
| **SM - GS** | **2,723** | **0,906** | **3,004** | **0,003** |
| T2 - T1 | -0,450 | 0,463 | -0,972 | 0,331 |
| **T3 - T1** | **-2,078** | **0,706** | **-2,943** | **0,003** |
| **T4 - T1** | **-2,830** | **0,807** | **-3,505** | **<0,001** |
| **T5 - T1** | **-1,922** | **0,632** | **-3,039** | **0,002** |
| **T6 - T1** | **-2,171** | **0,709** | **-3,063** | **0,002** |
| **T7 - T1** | **-1,975** | **0,726** | **-2,720** | **0,007** |
| T8 - T1 | -2,098 | 1,140 | -1,840 | 0,066 |

**Table 2.C. Post-hocs: Simple terms**

| IV | | Estimate | SE | Df | z.ratio | p.value |
|---|---|---|---|---|---|---|
| **Group** | **SM-GS** | **1,64** | **0,50** | **Inf** | **3,31** | **0,001** |
| **Time** | **T2 - T1** | **-1,17** | **0,38** | **Inf** | **-3,09** | **0,01** |
| | **T3 - T1** | **-2,61** | **0,51** | **Inf** | **-5,10** | **<0,001** |
| | **T4 - T1** | **-2,48** | **0,44** | **Inf** | **-5,66** | **<0,001** |
| | **T5 - T1** | **-2,18** | **0,42** | **Inf** | **-5,22** | **<0,001** |
| | **T6 - T1** | **-3,12** | **0,49** | **Inf** | **-6,41** | **<0,001** |

| | | | | | | |
|---|---|---|---|---|---|---|
| | **T7 - T1** | **-2,77** | **0,49** | **Inf** | **-5,65** | **<0,001** |
| | **T8 - T1** | **-2,24** | **0,60** | **Inf** | **-3,72** | **0,001** |

**Table 3 (#3): Perceived Sleep Depth ~ Group*Stage*Time + (1|sub) + (1|time asleep) in FAS – Fig.1.C-D**

**Table 3.A. Main effects**

| | Chisq | Df | Pr(>Chisq) |
|---|---|---|---|
| Group | 0,04 | 1 | 0,843096 |
| **Stage** | **19,71** | **1** | **<0,001** |
| **Time** | **17,56** | **7** | **<0,001** |
| **Group:Stage** | **5,54** | **1** | **0,02** |
| Group:Time | 1,65 | 7 | 0,20 |
| **Stage:Time** | **8,14** | **6** | **0,004** |
| Group:Stage:Time | 3,26 | 6 | 0,07 |

**Table 3.B. Estimates**

| | Estimate | Std Error | Df | T value | P value |
|---|---|---|---|---|---|
| **(Intercept)** | **3,27** | **0,16** | **68,96** | **20,87** | **<0,001** |
| SM - GS | -0,06 | 0,31 | 112,87 | -0,20 | 0,84 |
| **REM-NREM** | **1,24** | **0,28** | **614,56** | **4,44** | **<0,001** |
| T2 | 0,16 | 0,17 | 597,74 | 0,94 | 0,35 |
| **T3** | **0,50** | **0,17** | **594,00** | **2,91** | **0,00374** |
| **T4** | **0,75** | **0,16** | **593,49** | **4,69** | **<0,001** |
| **T5** | **0,90** | **0,17** | **594,92** | **5,25** | **<0,001** |
| **T6** | **0,60** | **0,17** | **596,09** | **3,45** | **<0,001** |
| **T7** | **0,75** | **0,18** | **594,06** | **4,12** | **<0,001** |
| T8 | -0,03 | 0,28 | 598,39 | -0,09 | 0,93 |

**Table 3.C. Post-hocs: Simple terms**

| IV | | Estimate | SE | Df | z.ratio | p.value |
|---|---|---|---|---|---|---|
| **Stage** | **NREM-REM** | **-0,46** | **0,08** | **Inf** | **-5,45** | **<0,001** |
| **Time** | **T2 - T1** | **-1,17** | **0,38** | **Inf** | **-3,09** | **0,01** |
| | **T3 - T1** | **-2,61** | **0,51** | **Inf** | **-5,10** | **<0,001** |
| | **T4 - T1** | **-2,48** | **0,44** | **Inf** | **-5,66** | **<0,001** |
| | **T5 - T1** | **-2,18** | **0,42** | **Inf** | **-5,22** | **<0,001** |
| | **T6 - T1** | **-3,12** | **0,49** | **Inf** | **-6,41** | **<0,001** |
| | **T7 - T1** | **-2,77** | **0,49** | **Inf** | **-5,65** | **<0,001** |
| | **T8 - T1** | **-2,24** | **0,60** | **Inf** | **-3,72** | **0,004** |

**Table 3.D. Post-hocs: Interaction terms**

| IV | contrast | estimate | SE | df | t.ratio | p.value |
|---|---|---|---|---|---|---|
| **Group: Stage** | GS_NREM - SM_NREM | 0,300 | 0,221 | 33,251 | 1,358 | 0,183 |
| | **GS_REM - SM_REM** | **0,696** | **0,243** | **49,983** | **2,865** | **0,006** |
| | **GS_NREM - GS_REM** | **-0,574** | **0,097** | **422,389** | **-5,901** | **<0,001** |
| | SM_NREM - SM_REM | -0,177 | 0,152 | 616,323 | -1,162 | 0,246 |
| **Stage :** | **NREM_2-REM_2** | **-0,516** | **0,251** | **609,289** | **-2,058** | **0,040** |
| | **NREM_3-REM_3** | **-0,663** | **0,219** | **577,651** | **-3,025** | **0,003** |

| | | | | | | |
|---|---|---|---|---|---|---|
| **Time** | NREM_4-REM_4 | -0,296 | 0,209 | 602,340 | -1,420 | 0,156 |
| | NREM_5-REM_5 | -0,372 | 0,190 | 600,169 | -1,961 | 0,050 |
| | NREM_6-REM_6 | -0,173 | 0,172 | 591,803 | -1,006 | 0,315 |
| | NREM_7-REM_7 | -0,100 | 0,193 | 598,914 | -0,520 | 0,604 |
| | **NREM_8-REM_8** | **-1,243** | **0,330** | **606,906** | **-3,769** | **<0,001** |
| | NREM_2-NREM_1 | 0,118 | 0,156 | 612,314 | 0,758 | 0,449 |
| | **NREM_3-NREM_1** | **0,536** | **0,151** | **607,955** | **3,555** | **<0,001** |
| | **NREM_4-NREM_1** | **0,787** | **0,143** | **610,565** | **5,485** | **<0,001** |
| | **NREM_5-NREM_1** | **0,833** | **0,152** | **613,448** | **5,470** | **<0,001** |
| | **NREM_6-NREM_1** | **0,566** | **0,151** | **613,387** | **3,746** | **<0,001** |
| | **NREM_7-NREM_1** | **0,702** | **0,161** | **608,787** | **4,375** | **<0,001** |
| | NREM_8-NREM_1 | -0,044 | 0,237 | 612,745 | -0,184 | 0,854 |
| | REM_3-REM_2 | 0,565 | 0,294 | 610,766 | 1,923 | 0,055 |
| | REM_4-REM_2 | 0,449 | 0,288 | 607,005 | 1,559 | 0,120 |
| | **REM_5-REM_2** | **0,571** | **0,276** | **612,136** | **2,071** | **0,039** |
| | REM_6-REM_2 | 0,105 | 0,263 | 611,671 | 0,398 | 0,691 |
| | REM_7-REM_2 | 0,169 | 0,269 | 609,202 | 0,627 | 0,531 |
| | REM_8-REM_2 | 0,565 | 0,339 | 614,380 | 1,669 | 0,096 |

**Table 4 (#1$^{bis}$): Perceived state ~ Group*Stage_all + (1|sub) + (1|time asleep) – Suppl Fig.1.A**

**Table 4.A Main effects**

| | Chisq | Df | P value |
|---|---|---|---|
| **Group** | **16,82** | **1** | **<0,001** |
| **Stage_all** | **12,37** | **2** | **0,002** |
| **Group :Stage_all** | **7,18** | **2** | **0,03** |

**Table 4.B. Estimates**

| | Estimate | Std Error | Z value | P value |
|---|---|---|---|---|
| **(Intercept)** | **-2,62** | **0,38** | **-6,83** | **<0,001** |
| **SM-GS** | **2,37** | **0,58** | **4,10** | **<0,001** |
| **N3-N2** | **0,73** | **0,34** | **2,11** | **0,03** |
| **REM-N2** | **-2,45** | **1,04** | **-2,36** | **0,02** |

**Table 4.C. Post-hocs: Simple Terms**

| IV | contrast | estimate | SE | t.ratio | P value |
|---|---|---|---|---|---|
| **Stage_all** | **N2 - REM** | **2,33** | **0,45** | **5,13** | **<0,001** |
| | **N3 - REM** | **2,46** | **0,48** | **5,17** | **<0,001** |
| | N2 – N3 | -0,13 | 0,26 | -0,50 | 0,62 |

**Table 4.D. Post-hocs: Interaction Terms**

| IV | contrast | estimate | SE | t.ratio | P value |
|---|---|---|---|---|---|
| **Group : Stage_all** | **SM_N2-GS_N2** | **2,37** | **0,58** | **4,10** | **<0,001** |
| | SM_N3-GS_N3 | 0,97 | 0,62 | 1,57 | 0,12 |
| | SM_REM-GS_REM | 2,39 | 1,21 | 1,98 | 0,05 |
| | SM_N3-SM_N2 | -0,67 | 0,41 | -1,65 | 0,10 |
| | **SM_REM-SM_N2** | **-2,43** | **0,51** | **-4,73** | **<0,001** |
| | **SM_N3-SM_REM** | **1,75** | **0,58** | **3,05** | **<0,001** |
| | **GS_N3-GS_N2** | **0,73** | **0,34** | **2,11** | **0,03** |
| | **GS_REM-GS_N2** | **-2,45** | **1,04** | **-2,36** | **0,02** |
| | **GS_N3-GS_REM** | **3,18** | **1,03** | **3,08** | **<0,001** |

**Table 5 (#3bis): Perceived sleep depth ~ Group*Stage_all*Time + (1|sub) + (1|time asleep) in FAS– Suppl Fig.1.B**

**Table 5.A Main effects**

| | Chisq | Df | P value |
|---|---|---|---|
| Group | 2,62 | 1 | 0,11 |
| **Stage_all** | **9,32** | **2** | **0,01** |
| **Time** | **32,72** | **7** | **0,009** |
| Group:Stage_all | 2,58 | 2 | 0,28 |
| Group:Time | 6,24 | 7 | 0,51 |
| Stage_all:Time | 20,12 | 13 | 0,09 |
| Group:Stage_all:Time | 9,08 | 12 | 0,70 |

**Table 5.B. Estimates**

| | Estimate | Std Error | Df | Z value | P value |
|---|---|---|---|---|---|
| **(Intercept)** | **3,17** | **0,26** | **347,11** | **12,34** | **<0,001** |
| SM-GS | -1,10 | 0,68 | 564,11 | -1,62 | 0,11 |
| N3-N2 | -0,02 | 0,27 | 585,95 | -0,06 | 0,95 |
| **REM-N2** | **1,24** | **0,41** | **580,95** | **3,05** | **0,002** |
| T2-T1 | -0,30 | 0,30 | 581,70 | -0,99 | 0,32 |
| T3-T1 | 0,23 | 0,28 | 581,04 | 0,82 | 0,41 |
| **T4-T1** | **0,69** | **0,27** | **580,66** | **2,57** | **0,01** |
| **T5-T1** | **0,75** | **0,27** | **583,92** | **2,74** | **0,01** |
| **T6-T1** | **0,55** | **0,28** | **583,76** | **1,97** | **0,049** |
| **T7-T1** | **0,59** | **0,28** | **581,31** | **2,10** | **0,04** |
| T8-T1 | -0,05 | 0,34 | 586,09 | -0,14 | 0,89 |

**Table 5.C. Post-hocs: Simple terms**

| | contrast | estimate | SE | df | t.ratio | P value |
|---|---|---|---|---|---|---|
| Stage_all | N2-N3 | -0,13 | 0,09 | 633,67 | -1,42 | 0,40 |
| | **N2-REM** | **-0,51** | **0,09** | **429,60** | **-5,56** | **<0,001** |
| | **N3-REM** | **-0,38** | **0,10** | **395,45** | **-3,92** | **<0,001** |
| Time | T2 - T1 | 0,21 | 0,15 | 618,94 | 1,43 | 0,69 |
| | **T3 - T1** | **0,70** | **0,14** | **615,68** | **4,84** | **<0,001** |
| | **T4 - T1** | **0,84** | **0,14** | **618,09** | **6,03** | **<0,001** |
| | **T5 - T1** | **0,95** | **0,14** | **622,74** | **6,64** | **<0,001** |
| | **T6 - T1** | **0,62** | **0,14** | **618,44** | **4,47** | **<0,001** |
| | **T7 - T1** | **0,73** | **0,15** | **620,96** | **4,99** | **<0,001** |
| | T8 - T1 | 0,47 | 0,20 | 622,10 | 2,34 | 0,13 |

**Table 6 (#2bis): Perceived State ~ Group*Time_since_first_sleep_onset + (1|sub) + (1|time asleep)**

**Table 6.A Main effects**

| | Chisq | Df | P value |
|---|---|---|---|
| **Group** | **8,72** | **1** | **0,003** |
| **Time_since_first_sleep_onset** | **31,68** | **7** | **<0,001** |
| Group : Time_since_first_sleep_onset | 3,98 | 7 | 0,78 |

**Table 6.B. Estimates**

| | Estimate | Std Error | Z value | P value |
|---|---|---|---|---|

| | | | | |
|---|---|---|---|---|
| **(Intercept)** | **-1,18** | **0,51** | **-2,29** | **0,02** |
| **groupP** | **2,72** | **0,92** | **2,95** | **0,03** |
| **T2-T1** | **-1,18** | **0,50** | **-2,35** | **0,02** |
| **T3-T1** | **-2,40** | **0,70** | **-3,44** | **<0,001** |
| **T4-T1** | **-2,37** | **0,69** | **-3,46** | **<0,001** |
| **T5-T1** | **-1,91** | **0,58** | **-3,30** | **<0,001** |
| **T6-T1** | **-2,72** | **0,81** | **-3,38** | **<0,001** |
| **T7-T1** | **-2,32** | **0,86** | **-2,70** | **0,01** |
| T8-T1 | -1,85 | 1,21 | -1,53 | 0,13 |

**Table 6.C. Post-hocs: Simple Terms**

| IV | contrast | estimate | SE | t.ratio | P value |
|---|---|---|---|---|---|
| **Group** | **SM-GS** | **1,85** | **0,61** | **3,02** | **0,003** |
| **Time_since_first_sleep_onset** | **T2-T1** | **-1,68** | **0,44** | **-3,86** | **<0,001** |
| | **T3-T1** | **-2,05** | **0,48** | **-4,31** | **<0,001** |
| | **T4-T1** | **-2,49** | **0,48** | **-5,18** | **<0,001** |
| | **T5-T1** | **-2,29** | **0,46** | **-4,94** | **<0,001** |
| | **T6-T1** | **-3,14** | **0,58** | **-5,42** | **<0,001** |
| | **T7-T1** | **-2,83** | **0,62** | **-4,59** | **<0,001** |
| | T8-T1 | -2,06 | 0,91 | -2,27 | 0,15 |

**Table 7 (#3$^{tris}$): Perceived Sleep Depth ~ Group * Stage * Time_since_first_sleep_onset + (1|sub) + (1|time asleep)**

**Table 7.A Main effects**

| | Chisq | Df | P value |
|---|---|---|---|
| **group** | **4,70** | **1** | **0,03** |
| **Stage_grouped** | **21,71** | **1** | **<0,001** |
| **time_hour_since_first_N1** | **50,33** | **7** | **<0,001** |
| **group:Stage_grouped** | **5,50** | **1** | **0,02** |
| group:time_hour_since_first_N1 | 9,84 | 7 | 0,20 |
| **Stage_grouped:time_hour_since_first_N1** | **13,05** | **6** | **0,04** |
| group:Stage_grouped:time_hour_since_first_N1 | 2,13 | 6 | 0,91 |

**Table 7.B. Estimates**

| | Estimate | Std Error | Df | Z value | P value |
|---|---|---|---|---|---|
| **(Intercept)** | **3,15** | **0,17** | **95,57** | **18,67** | **<0,001** |
| SM - GS | -0,14 | 0,35 | 166,57 | -0,40 | 0,69 |
| **REM - NREM** | **1,18** | **0,37** | **589,41** | **3,15** | **<0,001** |
| T2-T1 | 0,18 | 0,17 | 591,32 | 1,02 | 0,31 |
| **T3-T1** | **0,44** | **0,18** | **590,09** | **2,39** | **0,02** |
| **T4-T1** | **0,76** | **0,16** | **590,55** | **4,67** | **<0,001** |
| **T5-T1** | **0,87** | **0,18** | **592,64** | **4,92** | **<0,001** |
| **T6-T1** | **0,61** | **0,18** | **593,10** | **3,38** | **<0,001** |
| **T7-T1** | **0,78** | **0,18** | **592,51** | **4,32** | **<0,001** |
| T8-T1 | 0,07 | 0,27 | 592,40 | 0,26 | 0,79 |

**Table 7.C. Post-hocs: Simple terms**

| | contrast | estimate | SE | df | t.ratio | P value |
|---|---|---|---|---|---|---|
| **Group** | SM-GS | -0,39 | 0,21 | 29,19 | -1,85 | 0,08 |
| **Stage** | **NREM-REM** | **-0,47** | **0,08** | **294,18** | **-5,63** | **<0,001** |
| **Time** | T2 - T1 | 0,23 | 0,15 | 613,42 | 1,48 | 0,65 |
| | **T3 - T1** | **0,60** | **0,15** | **612,06** | **3,97** | **<0,001** |
| | **T4 - T1** | **0,85** | **0,14** | **612,87** | **5,91** | **<0,001** |
| | **T5 - T1** | **0,94** | **0,15** | **616,47** | **6,41** | **<0,001** |
| | **T6 - T1** | **0,63** | **0,15** | **617,80** | **4,32** | **<0,001** |
| | **T7 - T1** | **0,75** | **0,15** | **617,89** | **5,10** | **<0,001** |
| | **T8 - T1** | **0,72** | **0,20** | **617,83** | **3,64** | **0,002** |

#### II. Perceived sleep depth and conscious experiences

**Table 8 (#4): C.Exp (CE vs NE) ~ Group*Stage + (1|sub) + (1|time asleep) in FAS – Fig.2.C**

**Table 8.A. Main effects**

| | Chisq | Df | Pr(>Chisq) |
|---|---|---|---|
| Group | 0,93 | 1 | 0,33 |
| **Stage** | 12,60 | 1 | **<0,001** |
| Group:Stage | 0,05 | 1 | 0,82 |

**Table 8.B. Estimates**

| IV | Estimate | Std Error | Z value | P value |
|---|---|---|---|---|
| **(Intercept)** | **1,13** | **0,42** | **2,67** | **0,01** |
| SM-GS | -0,69 | 0,72 | -0,97 | 0,33 |
| **REM-NREM** | **1,34** | **0,38** | **3,55** | **<0,001** |

**Table 8.C. Post-hocs: Simple terms**

| | contrast | estimate | SE | df | t.ratio | P value |
|---|---|---|---|---|---|---|
| **Stage** | **NREM-REM** | **-1,39** | **0,30** | **Inf** | **-4,60** | **<0,001** |

**Table 9 (#5a): Thought-like ~ Group*Stage + (1|sub) + (1|time asleep) in FAS – Fig.2.D**

**Table 9.A. Main effects**

| | Chisq | Df | Pr(>Chisq) |
|---|---|---|---|
| **Group** | 9,01 | 1 | 0,003 |
| Stage | 0,22 | 1 | 0,64 |
| Group:Stage | 0,44 | 1 | 0,51 |

**Table 9.B. Estimates**

| | Estimate | Std Error | Df | T value | P value |
|---|---|---|---|---|---|
| **(Intercept)** | 2,14 | 0,16 | 28,02 | 13,13 | **<0,001** |
| **SM - GS** | 0,95 | 0,32 | 37,55 | 3,00 | 0,005 |
| REM - NREM | 0,08 | 0,18 | 286,51 | 0,47 | 0,64 |

**Table 9.C. Post-hocs: Simple terms**

| | contrast | estimate | SE | df | t.ratio | P value |
|---|---|---|---|---|---|---|
| **Group** | **SM-GS** | **1,06** | **0,28** | **28,29** | **3,75** | **0,001** |

**Table 10 (#5b): Perceptual ~ Group*Stage + (1|sub) + (1|time asleep) in FAS – Fig.2.D**
**Table 10.A. Main effects**

| | Chisq | Df | Pr(>Chisq) |
|---|---|---|---|
| Group | 1,55 | 1 | 0,21 |
| **Stage** | **10,58** | **1** | **0,001** |
| Group:Stage | 0,06 | 1 | 0,80 |

**Table 10.B. Estimates**

| | Estimate | Std Error | Df | T value | P value |
|---|---|---|---|---|---|
| **(Intercept)** | **2,27** | **0,22** | **27,67** | **10,17** | **<0,001** |
| SM-GS | 0,50 | 0,40 | 33,59 | 1,24 | 0,22 |
| **REM-NREM** | **0,50** | **0,15** | **217,59** | **3,25** | **0,001** |

**Table 10.C. Post-hocs: Simple terms**

| | contrast | estimate | SE | df | t.ratio | P value |
|---|---|---|---|---|---|---|
| **Stage** | **REM-NREM** | **0,50** | **0,13** | **285,65** | **3,93** | **0,0001** |

**Table 11 (#5c): Richness&Complexity ~ Group*Stage + (1|sub) + (1|time asleep) in FAS – Fig.2.D**

**Table 11.A. Main effects**

| | Chisq | Df | Pr(>Chisq) |
|---|---|---|---|
| Group | 0,01 | 1 | 0,94 |
| **Stage** | **32,01** | **1** | **<0,001** |
| Group:Stage | 0,11 | 1 | 0,74 |

**Table 11.B. Estimates**

| | Estimate | Std Error | Df | T value | P value |
|---|---|---|---|---|---|
| **(Intercept)** | 2,30 | 0,21 | 28,07 | 10,74 | **<0,001** |
| SM-GS | 0,03 | 0,39 | 34,07 | 0,07 | 0,94 |
| **REM-NREM** | 0,80 | 0,14 | 227,16 | 5,66 | **<0,001** |

**Table 11.C. Post-hocs: Simple terms**

| | contrast | estimate | SE | df | t.ratio | P value |
|---|---|---|---|---|---|---|
| **Stage** | **REM-NREM** | **0,76** | **0,12** | **285,06** | **6,45** | **<0,001** |

**Table 12 (#5d): Duration ~ Group*Stage + (1|sub) + (1|time asleep) in FAS – Fig.2.D**

**Table 12.A. Main effects**

| | Chisq | Df | Pr(>Chisq) |
|---|---|---|---|
| Group | 0,29 | 1 | 0,59 |
| **Stage** | **28,23** | **1** | **<0,001** |
| Group:Stage | 0,04 | 1 | 0,85 |

**Table 12.B. Estimates**

| | Estimate | Std Error | Df | T value | P value |
|---|---|---|---|---|---|
| **(Intercept)** | **2,07** | **0,18** | **28,27** | **11,57** | **<0,001** |
| SM-GS | -0,18 | 0,33 | 36,13 | -0,54 | 0,59 |

| | | | | | |
|---|---|---|---|---|---|
| REM-NREM | 0,73 | 0,14 | 207,19 | 5,31 | <0,001 |

**Table 12.C. Post-hocs: Simple terms**

| | contrast | estimate | SE | df | t.ratio | P value |
|---|---|---|---|---|---|---|
| Stage | REM-NREM | 0,71 | 0,12 | 287,84 | 6,14 | <0,001 |

**Table 13 (#5e): Control ~ Group*Stage + (1|sub) + (1|time asleep) in FAS – Fig.2.D**

**Table 13.A. Main effects**

| | Chisq | Df | Pr(>Chisq) |
|---|---|---|---|
| Group | 2,37 | 1 | 0,12 |
| Stage | 2,86 | 1 | 0,09 |
| Group:Stage | 0,01 | 1 | 0,91 |

**Table 13.B. Estimates**

| | Estimate | Std Error | Df | T value | P value |
|---|---|---|---|---|---|
| (Intercept) | 1,57 | 0,18 | 21,60 | 8,51 | <0,001 |
| SM-GS | 0,52 | 0,33 | 27,12 | 1,54 | 0,13 |
| REM-NREM | 0,22 | 0,13 | 270,04 | 1,69 | 0,09 |

**Table 14 (#5f): Lucidity ~ Group*Stage + (1|sub) + (1|time asleep) in FAS – Fig.2.D**

**Table 14.A. Main effects**

| | Chisq | Df | Pr(>Chisq) |
|---|---|---|---|
| Group | 0,32 | 1 | 0,57 |
| Stage | 5,33 | 1 | 0,02 |
| Group:Stage | 0,05 | 1 | 0,83 |

**Table 14.B. Estimates**

| | Estimate | Std Error | Df | T value | P value |
|---|---|---|---|---|---|
| (Intercept) | 2,19 | 0,22 | 25,77 | 9,76 | <0,001 |
| SM-GS | 0,23 | 0,40 | 32,33 | 0,57 | 0,57 |
| REM-NREM | -0,36 | 0,15 | 195,25 | -2,31 | 0,02 |

**Table 14.C. Post-hocs: Simple terms**

| | contrast | estimate | SE | df | t.ratio | P value |
|---|---|---|---|---|---|---|
| Stage | REM-NREM | -0,33 | 0,13 | 277,76 | -2,58 | 0,01 |

**Table 15 (#6): Perceived Sleep Depth ~ Group*Stage*CE + (1|sub) + (1|time asleep) in FAS**

**Table 15.A. Main effects**

| | Chisq | Df | Pr(>Chisq) |
|---|---|---|---|
| Group | 2,42 | 1 | 0,12 |
| Stage | 3,68 | 1 | 0,06 |
| C.Exp | 2,66 | 1 | 0,10 |
| Group:C.Exp | 2,06 | 1 | 0,15 |
| Stage:C.Exp | 0,08 | 1 | 0,77 |
| Group:Stage | 0,38 | 1 | 0,54 |

| | | | |
|---|---|---|---|
| **Group:Stage:C.Exp** | 0,24 | 1 | 0,63 |

**Table 15.B. Estimates**

| | Estimate | Std Error | Df | T value | P value |
|---|---|---|---|---|---|
| **(Intercept)** | 3,83 | 0,17 | 68,64 | 22,57 | **<0,001** |
| **SM - GS** | -0,49 | 0,32 | 92,11 | -1,56 | 0,12 |
| **REM - NREM** | 0,55 | 0,29 | 400,03 | 1,92 | 0,06 |
| **CE - NE** | -0,13 | 0,08 | 397,19 | -1,63 | 0,10 |

**Table 16 (#7a): Perceived Sleep Depth ~ Group*Stage*Thought-like + (1|sub) + (1|time asleep) in FAS**

**Table 16.A. Main effects**

| | Chisq | Df | Pr(>Chisq) |
|---|---|---|---|
| Group | 0,35 | 1 | 0,56 |
| **Stage** | **10,55** | **1** | **0,001** |
| Thought-like | 3,56 | 4 | 0,47 |
| Group:Thought-like | 1,63 | 4 | 0,80 |
| Stage:Thought-like | 2,44 | 4 | 0,66 |
| Group:Stage:Thought-like | 3,63 | 5 | 0,60 |

**Table 16.B. Estimates**

| | Estimate | Std Error | Df | T value | P value |
|---|---|---|---|---|---|
| **(Intercept)** | **3,60** | **0,17** | **51,40** | **21,74** | **<0,001** |
| **SM-GS** | -0,24 | 0,40 | 153,99 | -0,59 | 0,56 |
| **REM-NREM** | **0,67** | **0,21** | **255,71** | **3,25** | **0,001** |
| **Thought-like2** | -0,15 | 0,19 | 273,43 | -0,82 | 0,41 |
| **Thought-like3** | -0,01 | 0,23 | 263,38 | -0,05 | 0,96 |
| **Thought-like4** | 0,30 | 0,26 | 273,12 | 1,14 | 0,25 |
| **Thought-like5** | 0,34 | 0,35 | 275,90 | 0,96 | 0,34 |

**Table 16.C. Post-hocs: Simple terms**

| | contrast | estimate | SE | df | t.ratio | P value |
|---|---|---|---|---|---|---|
| **Stage** | **REM-NREM** | **0,46** | **0,08** | **340,02** | **5,45** | **<0,001** |

**Table 17 (#7b): Perceived Sleep Depth ~ Group*Stage*Perceptual + (1|sub) + (1|time asleep) in FAS**

**Table 17.A. Main effects**

| | Chisq | Df | Pr(>Chisq) |
|---|---|---|---|
| Group | 0,20 | 1 | 0,66 |
| **Stage** | **7,78** | **1** | **0,01** |
| **Perception** | **19,74** | **4** | **0,001** |
| Group:Perception | 1,64 | 4 | 0,80 |
| Stage:Perception | 1,70 | 4 | 0,79 |
| Group:Stage:Perception | 4,84 | 5 | 0,44 |

**Table 17.B. Estimates**

| | Estimate | Std Error | Df | T value | P value |
|---|---|---|---|---|---|
| **(Intercept)** | **3,28** | **0,17** | **63,85** | **19,58** | **<0,001** |
| SM-GS | 0,18 | 0,42 | 157,29 | 0,44 | 0,66 |

| | | | | | |
|---|---|---|---|---|---|
| **REM-NREM** | **0,70** | **0,25** | **254,59** | **2,79** | **0,01** |
| Perception2 | 0,31 | 0,19 | 265,83 | 1,67 | 0,10 |
| Perception3 | 0,35 | 0,21 | 276,78 | 1,62 | 0,11 |
| **Perception4** | **0,92** | **0,24** | **269,16** | **3,81** | **<0,001** |
| **Perception5** | **1,10** | **0,33** | **280,12** | **3,35** | **<0,001** |

**Table 17.C. Post-hocs: Simple terms**

| | contrast | estimate | SE | df | t.ratio | P value |
|---|---|---|---|---|---|---|
| **Stage** | **REM-NREM** | **0,46** | **0,08** | **340,02** | **5,45** | **<0,001** |
| **Perception** | **P2 – P1** | 0,29 | 0,15 | 284,85 | 1,97 | 0,19 |
| | **P3 – P1** | 0,30 | 0,16 | 294,46 | 1,89 | 0,22 |
| | **P4 – P1** | **0,84** | **0,18** | **295,89** | **4,73** | **<0,001** |
| | **P5 – P1** | **0,80** | **0,22** | **282,58** | **3,69** | **<0,001** |

**Table 18 (#7c): Perceived Sleep Depth ~ Group*Stage*Richness&Complexity + (1|sub) + (1|time asleep) in FAS**

**Table 18.A. Main effects**

| | Chisq | Df | Pr(>Chisq) |
|---|---|---|---|
| **Group** | 0,04 | 1 | 0,83 |
| **Stage** | 2,19 | 1 | 0,14 |
| **Richness&Complexity** | **17,86** | **4** | **0,001** |
| **Group:Richness&Complexity** | 8,58 | 4 | 0,07 |
| **Stage:Richness&Complexity** | 3,90 | 4 | 0,42 |
| **Group:Stage:Richness&Complexity** | 8,12 | 5 | 0,15 |

**Table 18.B. Estimates**

| | Estimate | Std Error | Df | T value | P value |
|---|---|---|---|---|---|
| **(Intercept)** | **3,48** | **0,17** | **64,80** | **19,96** | **<0,001** |
| **SM-GS** | -0,07 | 0,35 | 106,56 | -0,21 | 0,84 |
| **REM-NREM** | 0,47 | 0,32 | 259,16 | 1,48 | 0,14 |
| **Richness&Complexity2** | -0,13 | 0,19 | 274,09 | -0,65 | 0,52 |
| **Richness&Complexity3** | 0,11 | 0,22 | 281,79 | 0,49 | 0,62 |
| **Richness&Complexity4** | **0,87** | **0,26** | **280,27** | **3,27** | **0,001** |
| **Richness&Complexity5** | **0,79** | **0,38** | **257,24** | **2,06** | **0,04** |

**Table 18.C. Post-hocs: Simple terms**

| | contrast | estimate | SE | df | t.ratio | P value |
|---|---|---|---|---|---|---|
| **Richness&Complexity** | R2 – R1 | 0,02 | 0,16 | 290,51 | 0,12 | 1,00 |
| | R3 – R1 | 0,25 | 0,16 | 295,88 | 1,54 | 0,41 |
| | **R4 – R1** | **0,60** | **0,18** | **294,51** | **3,28** | **0,005** |
| | **R5 – R1** | **0,83** | **0,24** | **272,07** | **3,48** | **0,002** |

**Table 19 (#7d): Perceived Sleep Depth ~ Group*Stage*Duration + (1|sub) + (1|time asleep) in FAS**

**Table 19.A. Main effects**

| | Chisq | Df | Pr(>Chisq) |
|---|---|---|---|
| **Group** | 0,02 | 1 | 0,89 |
| **Stage** | 0,73 | 1 | 0,39 |
| **Duration** | 6,71 | 4 | 0,15 |

| | | | |
|---|---|---|---|
| **Group:Duration** | 2,01 | 4 | 0,73 |
| **Stage:Duration** | 2,35 | 4 | 0,67 |
| **Group:Stage:Duration** | 7,23 | 4 | 0,12 |

**Table 19.B. Estimates**

| | Estimate | Std Error | Df | T value | P value |
|---|---|---|---|---|---|
| **(Intercept)** | **3,47** | **0,17** | **57,05** | **20,25** | **<0,001** |
| **SM-GS** | -0,04 | 0,32 | 78,87 | -0,14 | 0,89 |
| **REM-NREM** | 0,28 | 0,33 | 277,40 | 0,85 | 0,40 |
| **Duration2** | 0,12 | 0,18 | 275,33 | 0,66 | 0,51 |
| **Duration3** | 0,26 | 0,23 | 276,14 | 1,13 | 0,26 |
| **Duration4** | **0,60** | **0,26** | **277,66** | **2,30** | **0,02** |
| **Duration5** | -0,45 | 0,64 | 265,46 | -0,69 | 0,49 |

**Table 20 (#7e): Perceived Sleep Depth ~ Group*Stage*Control + (1|sub) + (1|time asleep) in FAS**

**Table 20.A. Main effects**

| | Chisq | Df | Pr(>Chisq) |
|---|---|---|---|
| Group | 0,05 | 1 | 0,82 |
| **Stage** | **12,39** | **1** | **<0,001** |
| **Control** | **12,09** | **4** | **0,02** |
| Group:Control | 1,93 | 4 | 0,75 |
| Stage:Control | 6,39 | 4 | 0,17 |
| Group:Stage:Control | 7,64 | 5 | 0,18 |

**Table 20.B. Estimates**

| | Estimate | Std Error | Df | T value | P value |
|---|---|---|---|---|---|
| **(Intercept)** | **3,69** | **0,15** | **33,30** | **23,95** | **<0,001** |
| SM-GS | -0,07 | 0,31 | 51,19 | -0,23 | 0,82 |
| **REM-NREM** | **0,56** | **0,16** | **232,58** | **3,52** | **0,001** |
| Control2 | 0,09 | 0,21 | 269,97 | 0,40 | 0,69 |
| **Control3** | **-0,69** | **0,24** | **269,10** | **-2,82** | **0,01** |
| Control4 | -0,65 | 0,37 | 265,75 | -1,74 | 0,08 |
| Control5 | 0,57 | 0,89 | 265,40 | 0,64 | 0,52 |

**Table 20.C. Post-hocs: Simple terms**

| | contrast | estimate | SE | df | t.ratio | P value |
|---|---|---|---|---|---|---|
| **Stage** | **REM - NREM** | **0,46** | **0,08** | **340,02** | **5,45** | **<0,001** |
| Control | C2 – C1 | -0,04 | 0,16 | 286,57 | -0,28 | 1,00 |
| | C3 – C1 | -0,20 | 0,17 | 291,42 | -1,20 | 0,65 |
| | C4 – C1 | -0,50 | 0,28 | 290,56 | -1,75 | 0,29 |
| | C5 – C1 | 0,67 | 0,33 | 290,79 | 2,00 | 0,17 |

**Table 21 (#7f): Perceived Sleep Depth ~ Group*Stage*Lucidity + (1|sub) + (1|time asleep) in FAS**

**Table 21.A. Main effects**

| | Chisq | Df | Pr(>Chisq) |
|---|---|---|---|
| Group | 0,83 | 1 | 0,36 |
| **Stage** | **9,86** | **1** | **0,002** |
| Lucidity | 8,40 | 4 | 0,08 |

| | | | |
|---|---|---|---|
| Group:Lucidity | 4,01 | 4 | 0,40 |
| Stage:Lucidity | 6,65 | 4 | 0,16 |
| Group:Stage:Lucidity | 7,89 | 5 | 0,16 |

**Table 21.B. Estimates**

| | Estimate | Std Error | Df | T value | P value |
|---|---|---|---|---|---|
| **(Intercept)** | **3,87** | **0,17** | **44,37** | **22,51** | **<0,001** |
| SM-GS | -0,29 | 0,31 | 61,42 | -0,91 | 0,36 |
| **REM-NREM** | **0,54** | **0,17** | **235,87** | **3,14** | **0,002** |
| **Lucidity1** | **-0,51** | **0,20** | **266,82** | **-2,49** | **0,01** |
| Lucidity2 | -0,34 | 0,25 | 269,49 | -1,38 | 0,17 |
| **Lucidity3** | **-0,57** | **0,27** | **264,79** | **-2,16** | **0,03** |
| Lucidity4 | -0,23 | 0,33 | 254,91 | -0,68 | 0,50 |

**Table 21.C. Post-hocs: Simple terms**

| | contrast | estimate | SE | df | t.ratio | P value |
|---|---|---|---|---|---|---|
| **Stage** | **REM - NREM** | **0,46** | **0,08** | **340,02** | **5,45** | **<0,001** |

**Table 22 (#9a): Delta ~ Time + Group + Time:Group + (1|sub) + (1|time asleep) in NREM – Figure S.2**

**Table 22.A. Main effects**

| | Chisq | Df | Pr(>Chisq) |
|---|---|---|---|
| **Time** | **185,54** | **7** | **<0,001** |
| **Group** | **9,71** | **1** | **0,001** |
| **Time :Group** | **17,28** | **7** | **0,02** |

**Table 22.B. Estimates**

| | Estimate | Std Error | Df | T value | P value |
|---|---|---|---|---|---|
| **T2-T1** | **-2,45** | **0,71** | **535,41** | **-3,46** | **<0,001** |
| **T3-T1** | **-4,88** | **0,77** | **534,59** | **-6,38** | **<0,001** |
| **T4-T1** | **-6,12** | **0,70** | **534,64** | **-8,71** | **<0,001** |
| **T5-T1** | **-6,77** | **0,74** | **534,83** | **-9,18** | **<0,001** |
| **T6-T1** | **-7,13** | **0,76** | **535,45** | **-9,40** | **<0,001** |
| **T7-T1** | **-7,54** | **0,81** | **536,37** | **-9,35** | **<0,001** |
| **T8-T1** | **-8,73** | **1,27** | **535,90** | **-6,89** | **<0,001** |
| **SM-GS** | **-4,81** | **1,54** | **43,89** | **-3,12** | **0,003** |

**Table 22.C. Post-hocs: Simple terms**

| | contrast | estimate | SE | df | t.ratio | P value |
|---|---|---|---|---|---|---|
| **Time** | T2-T1 | -1,19 | 0,65 | 536,25 | -1,84 | 0,38 |
| | **T3-T1** | **-3,00** | **0,67** | **535,58** | **-4,49** | **<0,001** |
| | **T4-T1** | **-4,10** | **0,61** | **536,08** | **-6,68** | **<0,001** |
| | **T5-T1** | **-4,88** | **0,63** | **537,24** | **-7,79** | **<0,001** |
| | **T6-T1** | **-5,44** | **0,64** | **536,62** | **-8,44** | **<0,001** |
| | **T7-T1** | **-5,66** | **0,73** | **538,90** | **-7,74** | **<0,001** |
| | **T8-T1** | **-6,64** | **0,99** | **537,48** | **-6,73** | **<0,001** |
| Group | SM-GS | -1,64 | 1,37 | 28,85 | -1,19 | 0,24 |

**Table 22.D. Post-hocs: Interaction Terms**

| IV | contrast | estimate | SE | t.ratio | P value |
|---|---|---|---|---|---|
| Time : Group | **GS_T1-SM_T1** | **4,81** | **1,54** | **3,12** | **0,03** |

| | | | | | |
|---|---|---|---|---|---|
| | GS_T2-SM_T2 | 2,30 | 1,66 | 1,39 | 0,77 |
| | GS_T3-SM_T3 | 1,04 | 1,70 | 0,61 | 1,00 |
| | GS_T4-SM_T4 | 0,78 | 1,61 | 0,48 | 1,00 |
| | GS_T5-SM_T5 | 1,03 | 1,62 | 0,64 | 1,00 |
| | GS_T6-SM_T6 | 1,44 | 1,65 | 0,87 | 0,98 |
| | GS_T7-SM_T7 | 1,06 | 1,78 | 0,60 | 1,00 |
| | GS_T8-SM_T8 | 0,64 | 2,23 | 0,29 | 1,00 |

**Table 23 (#9b): Theta ~ Time + Group + Time:Group + (1|sub) + (1|time asleep) in NREM – Figure S.2**

**Table 23.A. Main effects**

| | Chisq | Df | Pr(>Chisq) |
|---|---|---|---|
| **Time** | **357,8054** | **7** | **<0,001** |
| **Group** | **4,34** | **1** | **0,037** |
| Time :Group | 11,91 | 7 | 0,10 |

**Table 23.B. Estimates**

| | Estimate | Std Error | Df | T value | P value |
|---|---|---|---|---|---|
| **T2-T1** | **-0,28** | **0,04** | **534,85** | **-6,49** | **<0,001** |
| **T3-T1** | **-0,50** | **0,05** | **534,59** | **-10,75** | **<0,001** |
| **T4-T1** | **-0,58** | **0,04** | **534,60** | **-13,63** | **<0,001** |
| **T5-T1** | **-0,61** | **0,04** | **534,67** | **-13,52** | **<0,001** |
| **T6-T1** | **-0,62** | **0,05** | **534,86** | **-13,54** | **<0,001** |
| **T7-T1** | **-0,62** | **0,05** | **535,15** | **-12,68** | **<0,001** |
| **T8-T1** | **-0,64** | **0,08** | **534,99** | **-8,26** | **<0,001** |
| **SM-GS** | **-0,32** | **0,15** | **33,15** | **-2,08** | **0,045** |

**Table 23.C. Post-hocs: Simple terms**

| | contrast | estimate | SE | df | t.ratio | P value |
|---|---|---|---|---|---|---|
| **Time** | T2-T1 | -0,21 | 0,04 | 534,43 | -5,45 | 5,51E-07 |
| | **T3-T1** | **-0,39** | **0,04** | **532,75** | **-9,71** | **<0,001** |
| | **T4-T1** | **-0,51** | **0,04** | **534,07** | **-13,58** | **<0,001** |
| | **T5-T1** | **-0,52** | **0,04** | **532,46** | **-13,63** | **<0,001** |
| | **T6-T1** | **-0,54** | **0,04** | **533,14** | **-13,86** | **<0,001** |
| | **T7-T1** | **-0,57** | **0,04** | **534,16** | **-12,94** | **<0,001** |
| | **T8-T1** | **-0,61** | **0,06** | **533,28** | **-10,20** | **<0,001** |
| Group | SM-GS | -0,18 | 0,15 | 28,26 | -1,24 | 0,23 |

**Table 24 (#9c): Alpha ~ Time + Group + Time:Group + (1|sub) + (1|time asleep) in NREM – Figure S.2**

**Table 24.A. Main effects**

| | Chisq | Df | Pr(>Chisq) |
|---|---|---|---|
| **Time** | **323,29** | **7** | **<0,001** |
| Group | 0,92 | 1 | 0,34 |
| Time :Group | 6,70 | 7 | 0,46 |

**Table 24.B. Estimates**

| | Estimate | Std Error | Df | T value | P value |
|---|---|---|---|---|---|

| | | | | | |
|---|---|---|---|---|---|
| **T2-T1** | **-0,15** | **0,03** | **534,57** | **-5,74** | **<0,001** |
| **T3-T1** | **-0,25** | **0,03** | **534,36** | **-8,45** | **<0,001** |
| **T4-T1** | **-0,29** | **0,03** | **534,37** | **-10,76** | **<0,001** |
| **T5-T1** | **-0,31** | **0,03** | **534,42** | **-11,17** | **<0,001** |
| **T6-T1** | **-0,32** | **0,03** | **534,58** | **-11,26** | **<0,001** |
| **T7-T1** | **-0,34** | **0,03** | **534,82** | **-11,04** | **<0,001** |
| **T8-T1** | **-0,31** | **0,05** | **534,69** | **-6,37** | **<0,001** |
| SM-GS | -0,15 | 0,11 | 32,07 | -1,39 | 0,17 |

**Table 24.C. Post-hocs: Simple terms**

| | contrast | estimate | SE | df | t.ratio | P value |
|---|---|---|---|---|---|---|
| **Time** | T2-T1 | **-0,12** | **0,02** | **531,78** | **-4,98** | **<0,001** |
| | **T3-T1** | **-0,20** | **0,03** | **532,85** | **-7,93** | **<0,001** |
| | **T4-T1** | **-0,26** | **0,02** | **533,40** | **-11,05** | **<0,001** |
| | **T5-T1** | **-0,27** | **0,02** | **534,86** | **-11,12** | **<0,001** |
| | **T6-T1** | **-0,30** | **0,02** | **534,30** | **-12,10** | **<0,001** |
| | **T7-T1** | **-0,33** | **0,03** | **535,20** | **-11,69** | **<0,001** |
| | **T8-T1** | **-0,32** | **0,04** | **530,54** | **-8,45** | **<0,001** |

**Table 25 (#9d): Sigma ~ Time + Group + Time:Group + (1|sub) + (1|time asleep) in NREM – Figure S.2**

**Table 25.A. Main effects**

| | Chisq | Df | Pr(>Chisq) |
|---|---|---|---|
| **Time** | **95,88** | **7** | **<0,001** |
| Group | 0,22 | 1 | 0,64 |
| **Time :Group** | **14,46** | **7** | **0,04** |

**Table 25.B. Estimates**

| | Estimate | Std Error | Df | T value | P value |
|---|---|---|---|---|---|
| **T2-T1** | -0,02 | 0,02 | 534,37 | -1,20 | 0,23 |
| **T3-T1** | **-0,07** | **0,02** | **532,21** | **-4,01** | <0,001 |
| **T4-T1** | **-0,08** | **0,02** | **532,14** | **-4,90** | <0,001 |
| **T5-T1** | **-0,06** | **0,02** | **528,21** | **-3,35** | **0,001** |
| **T6-T1** | **-0,05** | **0,02** | **534,72** | **-2,99** | **0,003** |
| **T7-T1** | **-0,07** | **0,02** | **528,70** | **-3,93** | <0,001 |
| **T8-T1** | **-0,08** | **0,03** | **534,40** | **-2,67** | **0,01** |
| **SM-GS** | 0,03 | 0,06 | 32,42 | 0,42 | 0,68 |

**Table 25.C. Post-hocs: Simple terms**

| | contrast | estimate | SE | df | t.ratio | P value |
|---|---|---|---|---|---|---|
| **Time** | **T2-T1** | **-0,05** | **0,01** | **534,50** | **-3,17** | **0,01** |
| | **T3-T1** | **-0,10** | **0,01** | **534,37** | **-6,75** | **<0,001** |
| | **T4-T1** | **-0,10** | **0,01** | **534,55** | **-7,27** | **<0,001** |
| | **T5-T1** | **-0,08** | **0,01** | **533,95** | **-5,91** | **<0,001** |
| | **T6-T1** | **-0,09** | **0,01** | **534,45** | **-6,51** | **<0,001** |
| | **T7-T1** | **-0,11** | **0,02** | **534,80** | **-6,76** | **<0,001** |
| | **T8-T1** | **-0,13** | **0,02** | **532,58** | **-5,97** | **<0,001** |
| **Time:Group** | H_T1-P_T1 | -0,03 | 0,06 | 32,43 | -0,42 | 1,00 |
| | H_T2-P_T2 | 0,03 | 0,06 | 35,60 | 0,46 | 1,00 |

| | | | | | | |
|---|---|---|---|---|---|---|
| | H_T3-P_T3 | 0,04 | 0,06 | 36,98 | 0,62 | 1,00 |
| | H_T4-P_T4 | 0,02 | 0,06 | 34,18 | 0,34 | 1,00 |
| | H_T5-P_T5 | 0,03 | 0,06 | 34,45 | 0,49 | 1,00 |
| | H_T6-P_T6 | 0,06 | 0,06 | 35,41 | 0,98 | 0,96 |
| | H_T7-P_T7 | 0,05 | 0,06 | 39,63 | 0,85 | 0,98 |
| | H_T8-P_T8 | 0,09 | 0,07 | 58,25 | 1,25 | 0,86 |

**Table 26 (#9e): Beta ~ Time + Group + Time:Group + (1|sub) + (1|time asleep) in NREM – Figure S.2**

**Table 26.A. Main effects**

| | Chisq | Df | Pr(>Chisq) |
|---|---|---|---|
| **Time** | 86,07 | 7 | **<0,001** |
| Group | 0,63 | 1 | 0,43 |
| Time :Group | 7,30 | 7 | 0,40 |

**Table 26.B. Estimates**

| | Estimate | Std Error | Df | T value | P value |
|---|---|---|---|---|---|
| T2-T1 | -1,22E-03 | 1,01E-03 | 534,75 | -1,22 | 0,22 |
| **T3-T1** | -3,63E-03 | 1,09E-03 | 534,50 | -3,34 | **<0,001** |
| **T4-T1** | -4,95E-03 | 9,97E-04 | 534,51 | -4,96 | **<0,001** |
| **T5-T1** | -4,39E-03 | 1,05E-03 | 534,58 | -4,20 | **<0,001** |
| **T6-T1** | -3,83E-03 | 1,08E-03 | 534,77 | -3,56 | **<0,001** |
| **T7-T1** | -4,76E-03 | 1,14E-03 | 535,04 | -4,16 | **<0,001** |
| **T8-T1** | -3,89E-03 | 1,80E-03 | 534,89 | -2,16 | 0,03 |
| SM-GS | -1,19E-03 | 3,67E-03 | 32,90 | -0,32 | 0,75 |

**Table 26.C. Post-hocs: Simple terms**

| | contrast | estimate | SE | df | t.ratio | P value |
|---|---|---|---|---|---|---|
| **Time** | T2-T1 | -1,25E-03 | 9,24E-04 | 531,88 | -1,36 | 0,74 |
| | **T3-T1** | **-5,24E-03** | **9,52E-04** | **532,92** | **-5,50** | **<0,001** |
| | **T4-T1** | **-6,00E-03** | **8,75E-04** | **533,50** | **-6,86** | **<0,001** |
| | **T5-T1** | **-5,45E-03** | **8,92E-04** | **535,01** | **-6,11** | **<0,001** |
| | **T6-T1** | **-5,16E-03** | **9,19E-04** | **534,42** | **-5,62** | **<0,001** |
| | **T7-T1** | **-6,37E-03** | **1,04E-03** | **535,42** | **-6,12** | **<0,001** |
| | T8-T1 | -3,18E-03 | 1,41E-03 | 530,70 | -2,26 | 0,16 |

**Table 27 (#9f): Gamma ~ Time + Group + Time:Group + (1|sub) + (1|time asleep) in NREM – Figure S.2**

**Table 27.A. Main effects**

| | Chisq | Df | Pr(>Chisq) |
|---|---|---|---|
| **Time** | **32,82** | **7** | **<0,001** |
| Group | 1,24 | 1 | 0,27 |
| Time :Group | 10,94 | 7 | 0,14 |

**Table 27.B. Estimates**

| | Estimate | Std Error | Df | T value | P value |
|---|---|---|---|---|---|
| **T2-T1** | **0,01** | **8,29E-04** | **37,41** | **13,68** | **<0,001** |
| T3-T1 | -6,70E-04 | 4,99E-04 | 535,52 | -1,34 | 0,18 |

| | | | | | |
|---|---|---|---|---|---|
| T4-T1 | -4,26E-04 | 5,39E-04 | 535,10 | -0,79 | 0,43 |
| **T5-T1** | **-1,75E-03** | **4,95E-04** | **535,12** | **-3,53** | **<0,001** |
| **T6-T1** | **-1,25E-03** | **5,19E-04** | **535,23** | **-2,41** | **0,02** |
| T7-T1 | -8,33E-04 | 5,34E-04 | 535,54 | -1,56 | 0,12 |
| T8-T1 | -9,45E-04 | 5,67E-04 | 536,01 | -1,67 | 0,10 |
| SM-GS | -1,19E-03 | 8,93E-04 | 535,75 | -1,33 | 0,18 |

**Table 27.C. Post-hocs: Simple terms**

| | contrast | estimate | SE | df | t.ratio | P value |
|---|---|---|---|---|---|---|
| **Time** | T2-T1 | -3,51E-04 | 4,58E-04 | 532,37 | -0,77 | 0,98 |
| | **T3-T1** | **-1,36E-03** | **4,72E-04** | **533,26** | **-2,88** | **0,03** |
| | **T4-T1** | **-2,16E-03** | **4,34E-04** | **533,94** | **-4,97** | **<0,001** |
| | **T5-T1** | **-1,69E-03** | **4,43E-04** | **535,72** | **-3,82** | **0,001** |
| | **T6-T1** | **-1,63E-03** | **4,56E-04** | **535,00** | **-3,57** | **0,002** |
| | **T7-T1** | **-1,88E-03** | **5,16E-04** | **536,48** | **-3,65** | **0,002** |
| | T8-T1 | -1,34E-03 | 6,99E-04 | 531,44 | -1,91 | 0,33 |
